# Schlafen14 Regulates Gene Expression Depending on Codon Usage

**DOI:** 10.1101/2023.03.19.533350

**Authors:** Carlos Valenzuela, Sergio Saucedo, Manuel Llano

## Abstract

Schlafen (SLFN) is a family of proteins upregulated by type I interferons with a regulatory role in translation. Intriguingly, SLFN14 associates with the ribosome and can degrade rRNA, tRNA, and mRNA *in vitro*, but a role in translation is still unknown. Ribosomes are important regulatory hubs during translation elongation of mRNAs rich in rare codons. Therefore, we evaluated the potential role of SLFN14 in the expression of mRNAs enriched in rare codons, using HIV-1 genes as a model. We found that SLFN14 regulates in a variety of cell types, including primary immune cells, the expression of HIV-1 and non-viral genes based on their codon adaptation index, a measurement of the synonymous codon usage bias; consequently, SLFN14 inhibited the replication of HIV-1. The potent inhibitory effect of SLFN14 on the expression of the rare codon-rich transcript HIV-1 Gag was minimized by codon optimization. Mechanistically, we found that the endoribonuclease activity of SLFN14 is required, and that ribosomal, but not messenger, RNA degradation is associated with the transcript selectivity of SLFN14. Therefore, we propose that SLFN14 impairs the expression of transcripts rich in rare codons, in a catalytic-dependent manner.

## INTRODUCTION

Codon usage is one of the factors influencing the amount of polypeptide produced per mRNA in bacteria and eukaryotic cells and tRNA abundance seems to be a major factor in this process [1–12]. The relative deficiency of charged tRNAs, associated with the translation of transcripts rich in rare codons, causes ribosome pausing reducing the translation elongation rate and leading to ribosome stalling [7, 13–16]. This event decreases global protein synthesis by premature termination and degradation of the truncated protein through the ribosome-associated protein quality control [12, 17, 18], degradation of the associated mRNAs through the No-go decay mechanism [19, 20] [21] [22], rRNA degradation through the nonfunctional 18S rRNA decay pathway, or by inhibit translation initiation via the Integrated Stress Response [23].

The tRNA repertoire is modified during stress [24–31] including viral infection [32–36]. The innate immune system has evolved mechanisms to regulate protein expression based on rare codon usage bias through the effect of the type I interferon (IFN)-induced protein SLFN11. By tRNA degradation, SLFN11 negatively regulates expression of proteins encoded by transcripts with bias towards rare codons [32, 33, 37, 38]. SLFN13 also degrades tRNAs [39], whereas SLFN5 and SLFN2 bind to tRNA without degrading them, and through this interaction, SLFN2 protects stress-induced, angiogenin-mediated tRNA degradation [28, 40].

Another member of the SLFN family, SLFN14, was discovered as a stalled ribosome-associated endoribonuclease, and the purified protein was shown to degrade ribosomes, tRNAs, and mRNA *in vitro* [41–43]. Furthermore, SLFN14 was found to degrade LINE-1 mRNA [44] in cells. SLFN14 was also reported to impair the expression of two varicella zoster virus proteins, immediately early 62 and glycoprotein E [44, 45], and the nucleoprotein from influenza virus [45]. Intriguingly, all these viral proteins are encoded by transcripts enriched in rare codons that exhibit low codon adaptation indexes (CAI), a measurement of the synonymous codon usage bias [46]. For example, varicella zoster virus proteins, immediately early 62 and glycoprotein E have a CAI of 0.72 and 0.7, respectively, and the nucleoprotein from influenza virus has a CAI of 0.753. Therefore, we postulated that SLFN14 could inhibit translation of transcripts with bias toward rare codons in response to constraints that these transcripts likely encounter during translation elongation, i.e., ribosome stalling. To evaluate our hypothesis, we selected as a model HIV-1 genome which is rich in adenine nucleotides determining a codon usage bias that importantly differ from that of the host [47–49].

Our data indicate for the first time that SLFN14 potently inhibits expression of transcripts rich in rare codons, i.e., HIV-1 Gag and firefly luciferase; but no of those poor in rare codons, i.e., codon optimized HIV-1 Gag, HIV-1 Tat, cyan, monomeric (m) cherry, and green fluorescent proteins, and human CD4. As a consequence, HIV-1 replication is inhibited by SLFN14. The anti-HIV-1 effect was observed in primary monocytes and CD4+ T lymphocyte, and in cancer ( SUP-T1) and transformed (HEK293T) cell lines. This activity is type I IFN-independent, but requires the endoribonuclease function of SLFN14. Expression of SLFN14 was associated with ribosome degradation in cells co-expressing wild type but not codon optimized HIV-1 Gag mRNA. In sum, ours results indicate a novel function of SLFN14 in restricting expression of transcripts rich in rare codons.

## RESULTS

### Human and mouse SLFN14 preferentially impairs the expression of proteins enriched in rare codons

To evaluate the effect of SLFN14 on the expression of genes enriched in rare codons we used HIV-1 as a model. Advantageously, HIV-1 gene expression can be studied with plasmids encoding this virus or its individual proteins, or using the copy of the viral genome integrated into the host chromosome, whose expression is regulated as any cellular gene is. Furthermore, HIV-1 has open reading frames with different codon usage that are included in a common transcript produced from the viral promoter. Moreover, different reporter genes can be efficiently inserted into the HIV-1 genome. To study HIV-1 protein expression, we used the methodology described before to determine the anti-HIV-1 activity of SLFN11 [32, 33], SLFN13 [39], and N4BP1 [50]. This method has been also extensively used to evaluate the production phase of the HIV-1 life cycle. In this experimental strategy, HEK293T cells are co-transfected with plasmids expressing the proteins being evaluated and a plasmid encoding HIV-1. Because HEK293T cells lack detectable expression of SLFN14 mRNA [41] and proteins (see figure 3A below), these cells are suitable to study SLFN14.

The HIV-1 reporter [51] that we used to analyze the activity of SLFN14 expresses from the viral promoter a transcript that contains open reading frames with synonymous codon-usage that greatly differ from the host, such as Gag (CAI of 0.56). A fraction of this transcript experiences splicing, generating other transcripts containing open reading frames whose synonymous codon-usage resemble the host preferences, as for example Tat (CAI of 0.761), and a reporter gene (eGFP) that is codon optimized for human cells (CAI of 0.962). Transcription of this precursor mRNA is Tat-dependent. Therefore, this reporter offers the possibility to evaluate the effect of SLFN14 on the expression of transcripts with three different CAIs.

BLAST analysis[52] indicated that the primary structure of human and mouse SLFN14 proteins is 70% identical, representing the highest evolutionary conservation between human SLFNs and their mouse orthologs in the SLFN group III [53]. Therefore, we hypothesized that mouse and human SLFN14 will share relevant biological functions, and tested this prediction in our experiments.

HEK293T cells were co-transfected with the plasmid expressing this HIV-1 reporter [51] and a plasmid expressing cyan fluorescence protein (CFP), together with either a plasmid empty (control) or plasmids expressing mouse or human SLFN14. Seventy-two hours later expression of CFP and eGFP, and HIV-1 p24 (a limited proteolysis product of Gag) was analyzed by flow cytometry and ELISA, respectively. HIV-1 p24 values were normalized to the % of CFP+ cells to account for transfection efficiency. In addition, levels of SLFN14 in these cells was determined by immunoblot analysis.

Data in Fig. 1A-I showed that cells expressing either mouse or human SLFN14 produced ∼86% less HIV-1 p24 than control cells. Despite this severe reduction in HIV-1 p24, the eGFP Mean Fluorescence Intensity (MFI), an estimate the number eGFP molecules per cell, was similar in cells expressing or no SLFN14 proteins (Fig. 1A-II). SLFN14 levels were verified in these cells by immunoblot (Fig. 1A-III). Because the transcript encoding eGFP derives from the messenger encoding Gag, preservation of eGFP expression indicated that SLFN14 impaired Gag production at a post-transcriptional step. Furthermore, since in this model HIV-1 transcription depends of Tat, the levels of this protein were not diminished by SLFN14 proteins, at least importantly. Therefore, SLFN14 proteins selectively impaired the expression of Gag, the lower CAI protein in the group analyzed.

**Fig. 1.**
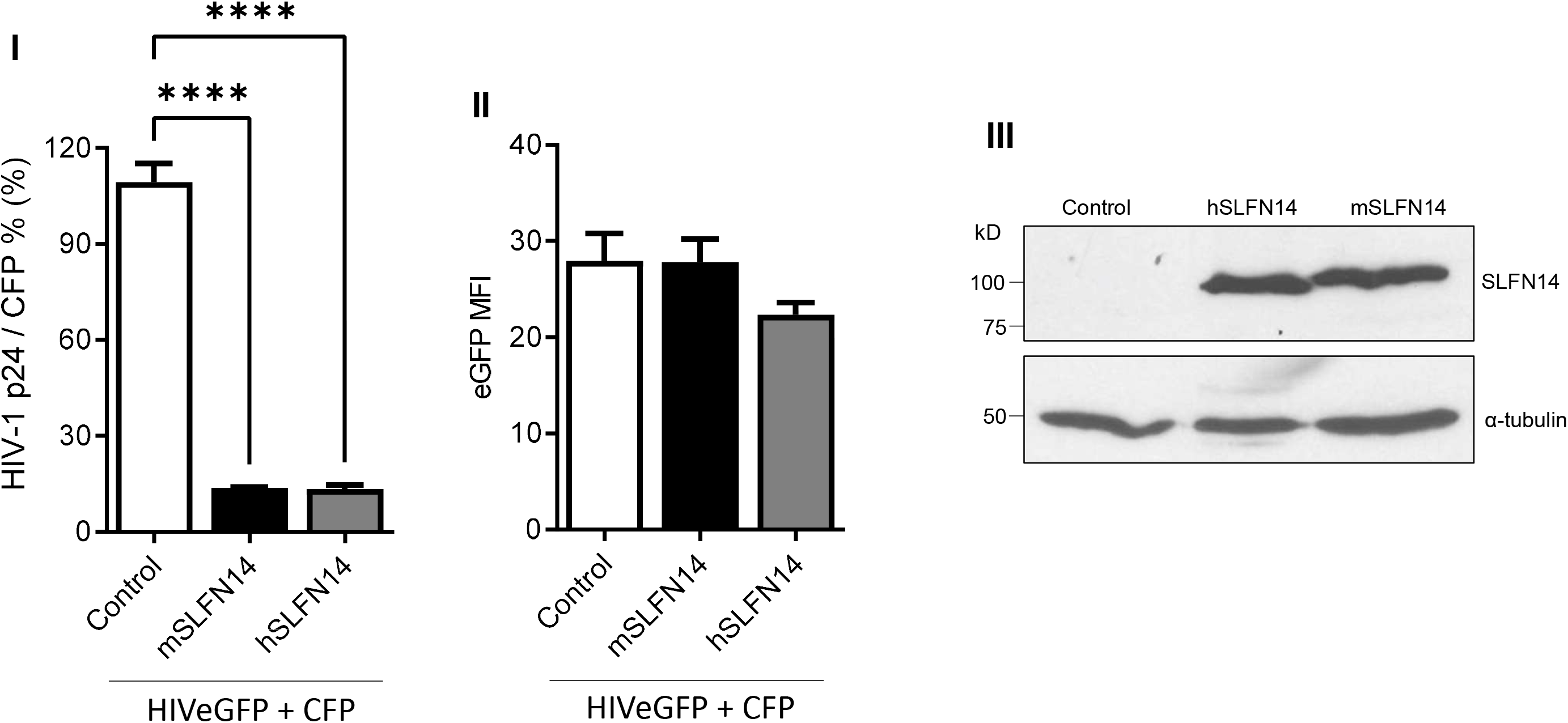
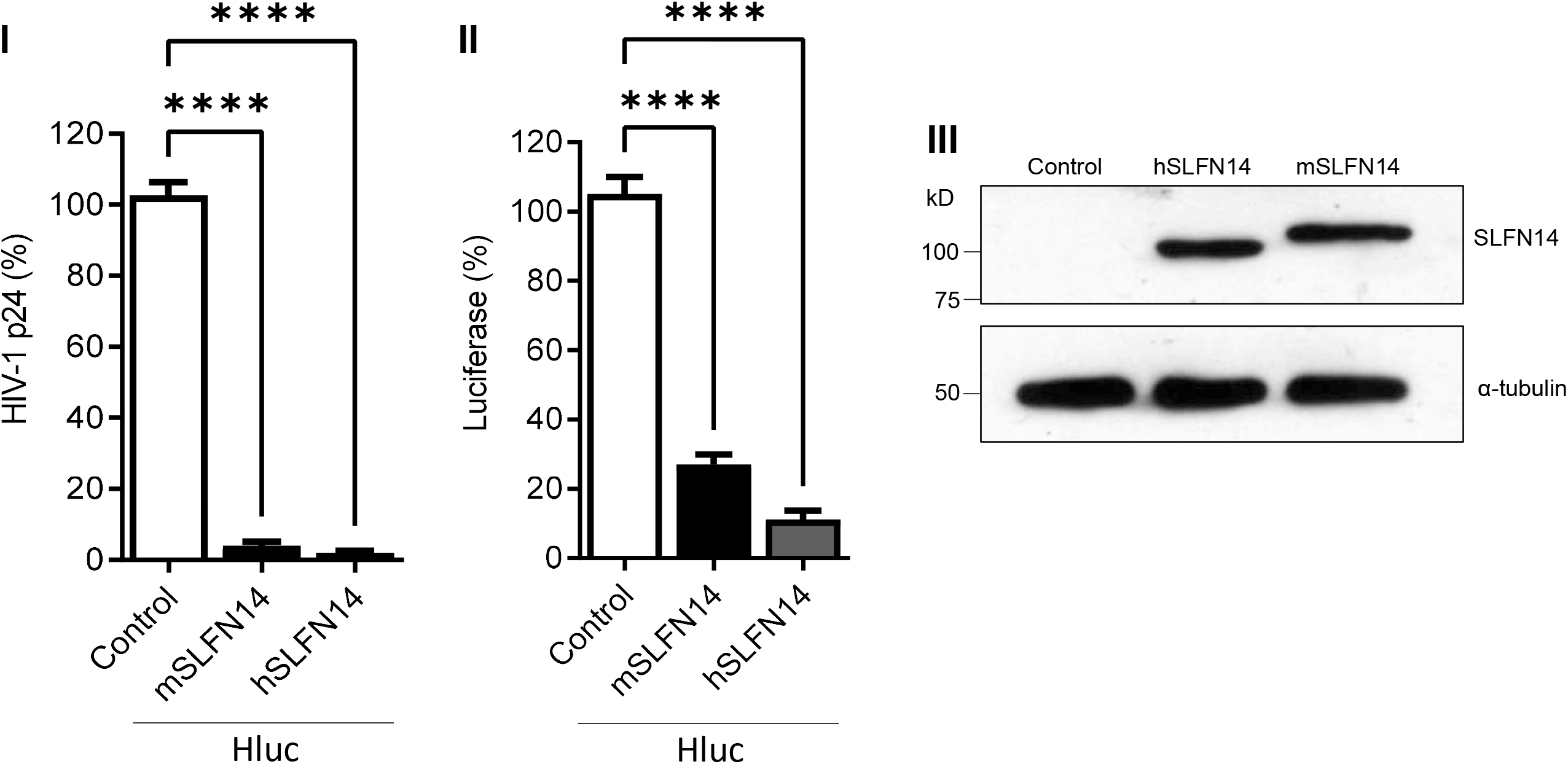
Effect of SLFN14 on HIV-1-driven protein expression. **(A)**. HEK293T cells that were co-transfected with a plasmid encoding an HIV-1 virus expressing eGFP and a plasmid expressing CFP together and either an empty plasmid (control cells) or a plasmid expressing human (h) or mouse (m) SLFN14. **(I)** HIV-1 p24 levels were normalized for transfection efficiency (% of CFP+ cells) and expressed as % of control cells. **(II)** eGFP MFI levels and **(III)** SLFN14 protein expression in cells represented in panel **(I)**. In the immunoblot **(III)** an anti-FLAG antibody was used to detect SLFN14 and α-tubulin levels were determined as a loading control. Data correspond to a triplicate experiment and are representative of five independent experiments. **(B)** HIV-1 p24 **(I)** and luciferase levels **(II)** expressed as % of control cells in HEK293T cells that were co-transfected with a plasmid encoding an HIV-1 virus expressing firefly luciferase (Hluc) and either an empty plasmid (control cells) or a plasmid expressing human (h) or mouse (m) SLFN14. **(III)** Expression of SLFN14 in these cells was detected by immunoblot as described in Fig. 1A-III. Data correspond to a triplicate experiment and are representative of more than twenty independent experiments performed along several months. Statistically significance in **(A)** and **(B)** was calculated with one-way ANOVA and Dunnett *post hoc* tests. **** *P* ≤ 0.0001. Data showing not statistically significant differences (NS P > 0.05) were not indicated in any of the figures of this work.

Next, we used a different HIV-1 reporter [54] that expresses Tat-dependent, viral promoter-driven firefly luciferase (CAI of 0.71), instead of eGFP. As before, HEK293T cells were co-transfected with an empty plasmid (control) or plasmids expressing human or mouse SLFN14, and a plasmid encoding the luciferase HIV-1 reporter, and analyzed seventy-two hrs later. Again, cells expressing mouse and human SLFN14 produced ∼97% less HIV-1 p24 than the control cells (Fig. 1B-I); whereas, luciferase expression was reduced by ∼73% and ∼89% by mouse and human SLFN14, respectively (Fig. 1B-II). SLFN14 levels were verified in these cells by immunoblot (Fig. 1A-III). These findings further demonstrated that SLFN14 impairs the expression of viral and non-viral proteins with low CAIs.

We also determined the effect of SLFN14 proteins on luciferase levels using a simpler expression system. HEK293T cells were co-transfected with the empty plasmid or plasmids expressing SLFN14 proteins, and a plasmid expressing firefly luciferase from a CMV enhancer/promoter (pCI Luc), and luciferase activity was measured three days after transfection. In these experiments, SLFN14-transfected cells produced ∼79 % (mouse) and ∼53 % (human) less luciferase than the control cells (Fig. 2A-I), indicating an HIV-1-independent effect of SLFN14 proteins on the expression of luciferase. SLFN14 levels were verified in these cells by immunoblot (Fig. 2A-II).

**Fig. 2.**
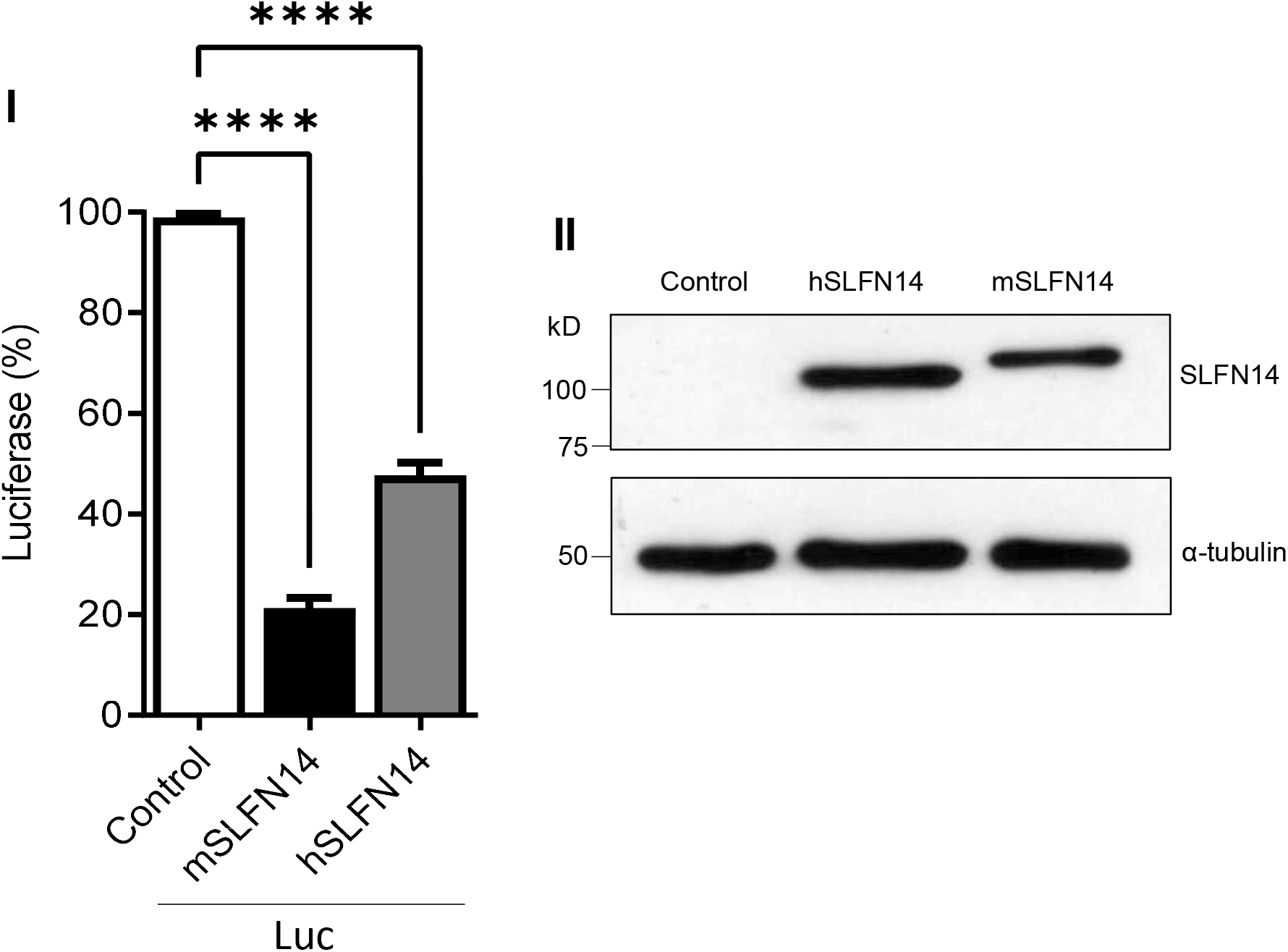
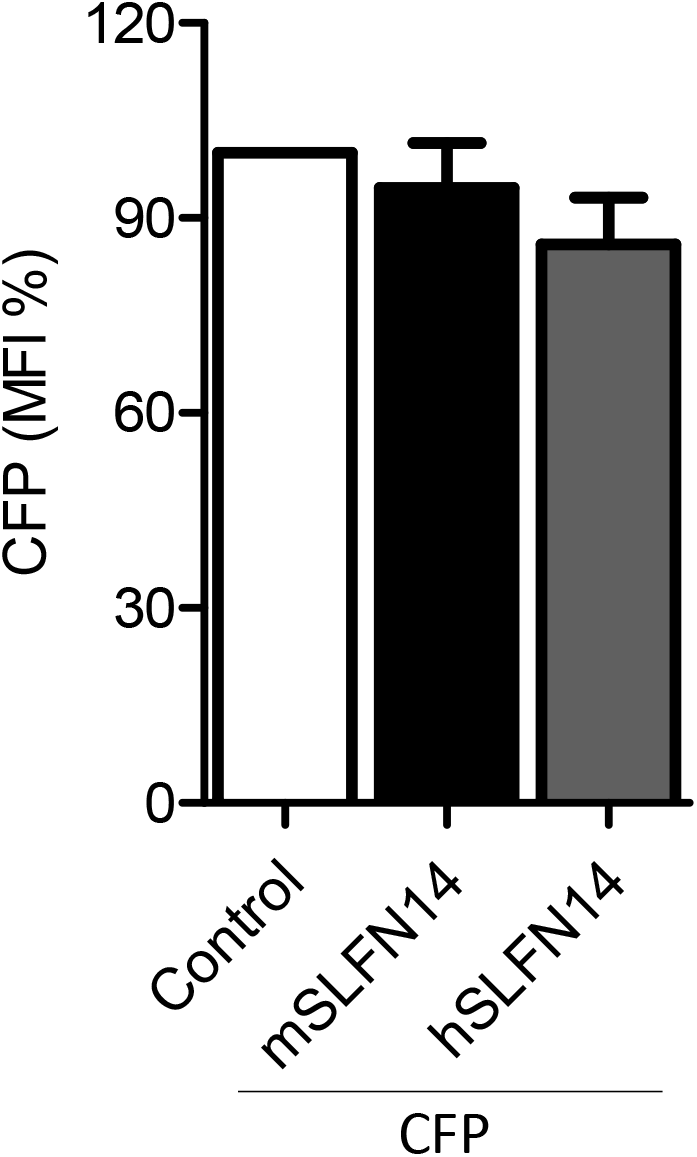
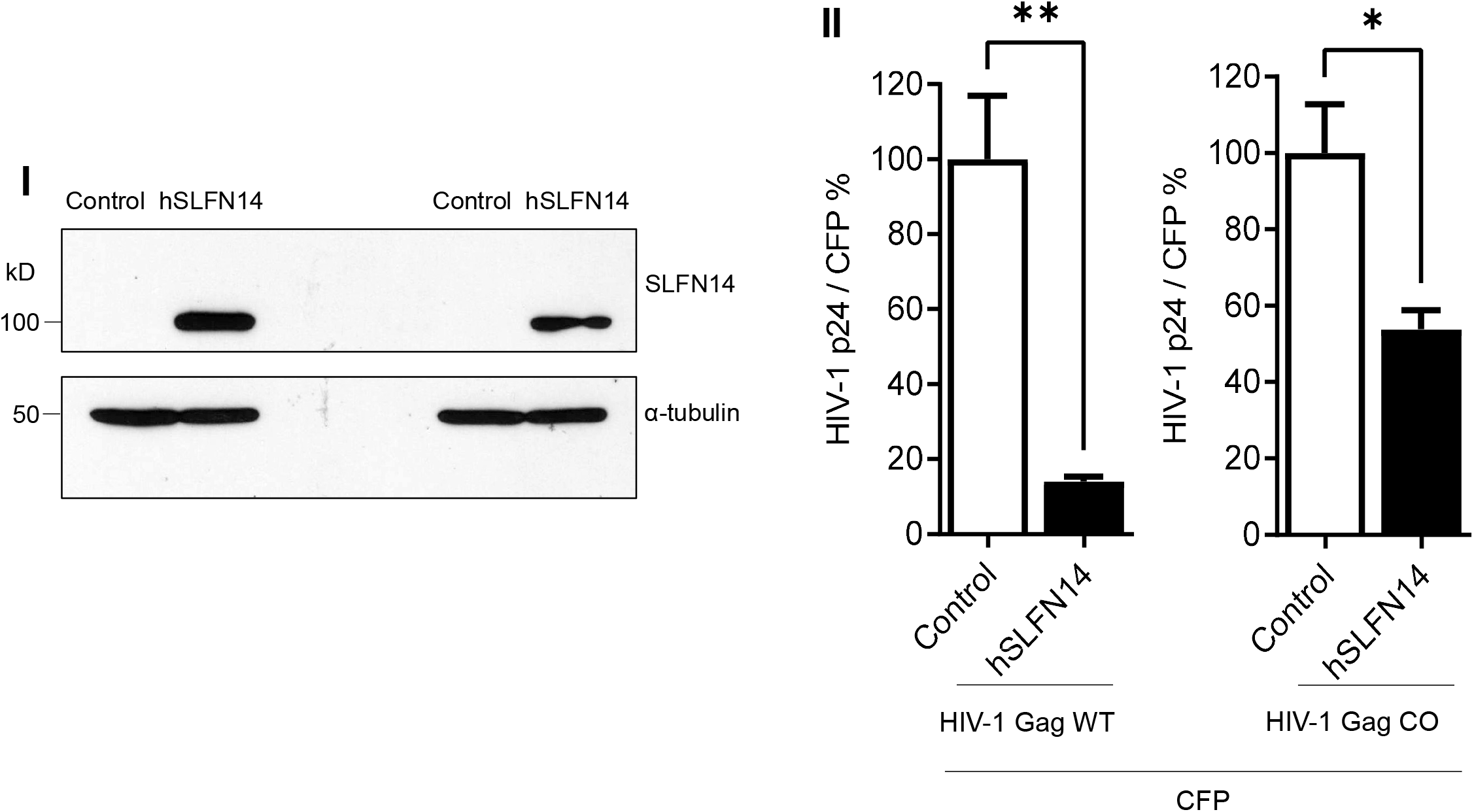
Effect of SLFN14 on the expression of transcripts with different codon usage. HEK293T cells were co-transfected with a plasmid expressing firefly luciferase or CFP and either an empty plasmid (control cells) or a plasmid expressing human (h) or mouse (m) SLFN14. Luciferase activity **(A-I)** and CFP MIF **(B)** were expressed as % of control cells. **(A-III)** Expression of SLFN14 in these cells was detected as described in Fig. 1A-III. Statistical significance was determined by one-way ANOVA and Dunnett *post hoc* tests. **** *P* ≤ 0.0001. Data correspond to a triplicate experiment and are representative of five **(A-I)** or six **(B)** independent experiments. **(C)** Effect of SLFN14 on HIV-1 Gag wild type (WT) and codon optimized (CO) expression. HEK293T cells were co-transfected with a plasmid expressing CFP and plasmids expressing either HIV-1 Gag WT or CO together with either an empty plasmid or a SLFN14 expression plasmid. **(I)** Expression of SLFN14 in these cells was detected as described in Fig. 1A-III. **(II)** HIV-1 p24 was normalized for transfection efficiency (% of CFP+ cells) and expressed as % of control cells. Statistically significant differences were calculated with repeated-measures using two-tailed, two sample t test. ** *P* ≤ 0.01 and * *P* ≤ 0.05. Data correspond to a triplicate experiment and are representative of eight independent experiments.

The effect of SLFN14 on the expression of a codon optimized messenger in a simpler expression system was also evaluated. HEK293T cells were co-transfected with a plasmid expressing CFP (CAI of 0.96) and plasmids expressing mouse or human SLFN14 or not (empty, control). Transfected cells were evaluated by flow cytometry three days later. As data in Fig. 2B indicate, expression of mouse or human SLFN14 did not reduce CFP MFI, in comparison to the control cells, indicating again that SLFN14 does not affect expression of codon optimized messengers.

To further evaluate a potential function of SLFN14 in codon usage-based control of gene expression, we determined the effect of SLFN14 on the expression of HIV-1 Gag wild type and codon optimized (CAI of 0.99). These Gag open reading frames are under the transcriptional control of the CMV enhancer/promoter. HEK293T cells were co-transfected with plasmids expressing CFP and either empty plasmid or SLFN14 expression plasmid together with plasmids expressing wild type or codon optimized Gag. HIV-1 p24 and CFP levels were measured 72 hrs after transfection by ELISA and flow cytometry, respectively, and HIV-1 p24 levels were normalized to the transfection efficiency (% of CFP+ cells). In these experiments, despite that similar levels of SLFN14 were achieved in cells expressing wild type and codon optimized Gag (Fig. 2C-I), SLFN14 impaired by ∼10 folds the expression of wild type Gag and by only ∼1.8 folds the expression of codon optimized Gag (Fig. 2C-II), indicating that SLFN14 preferentially impairs expression of transcripts enriched in rare codons.

### Expression of SLFN14 in cells of the immune system

Since SLFN14 potently impairs HIV-1 Gag expression, we predicted that this protein should not be very abundant in cells naturally permissive to HIV-1. Therefore, we characterized the expression of endogenous SLFN14 in cells of the immune system by immunoblot. In this analysis we used the only two anti-SLFN14 antibodies commercially available. These antibodies recognize the N-terminal (residues 100 -250) and the C-terminal (amino acids 730 – 780) regions of SLFN14. As control, we analyzed HEK293T since SLFN14 mRNA was reported undetectable in these cells [41]. HEK293T cells were transfected with an empty plasmid or a plasmid expressing human SLFN14 and seventy-two hrs later analyzed by immunoblot. As expected, no endogenous SLFN14 was detected in these cells with any of the anti-SLFN14 antibodies evaluated although both recognized the exogenous SLFN14 (Fig. 3A).

**Figure 3.**
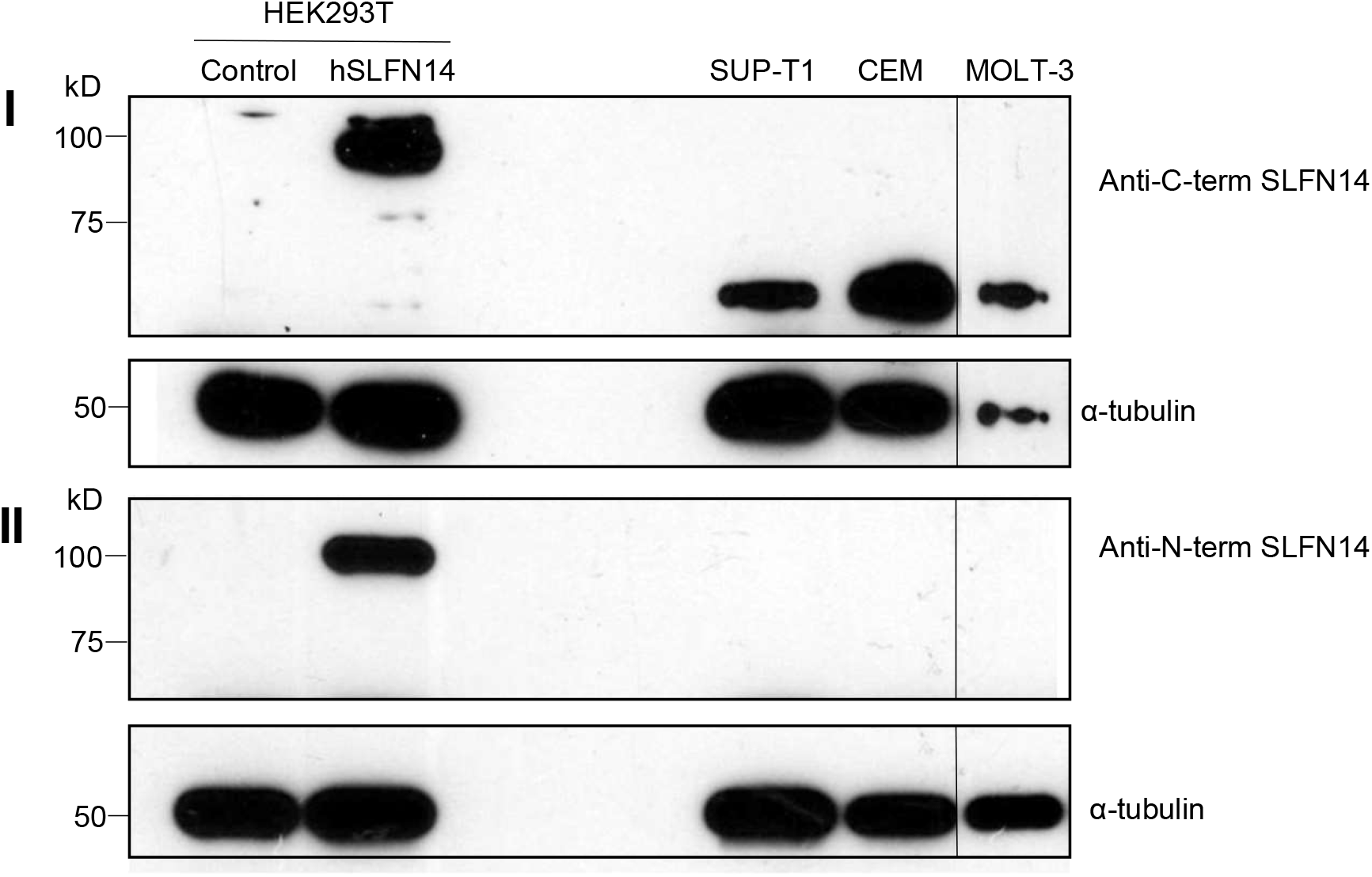
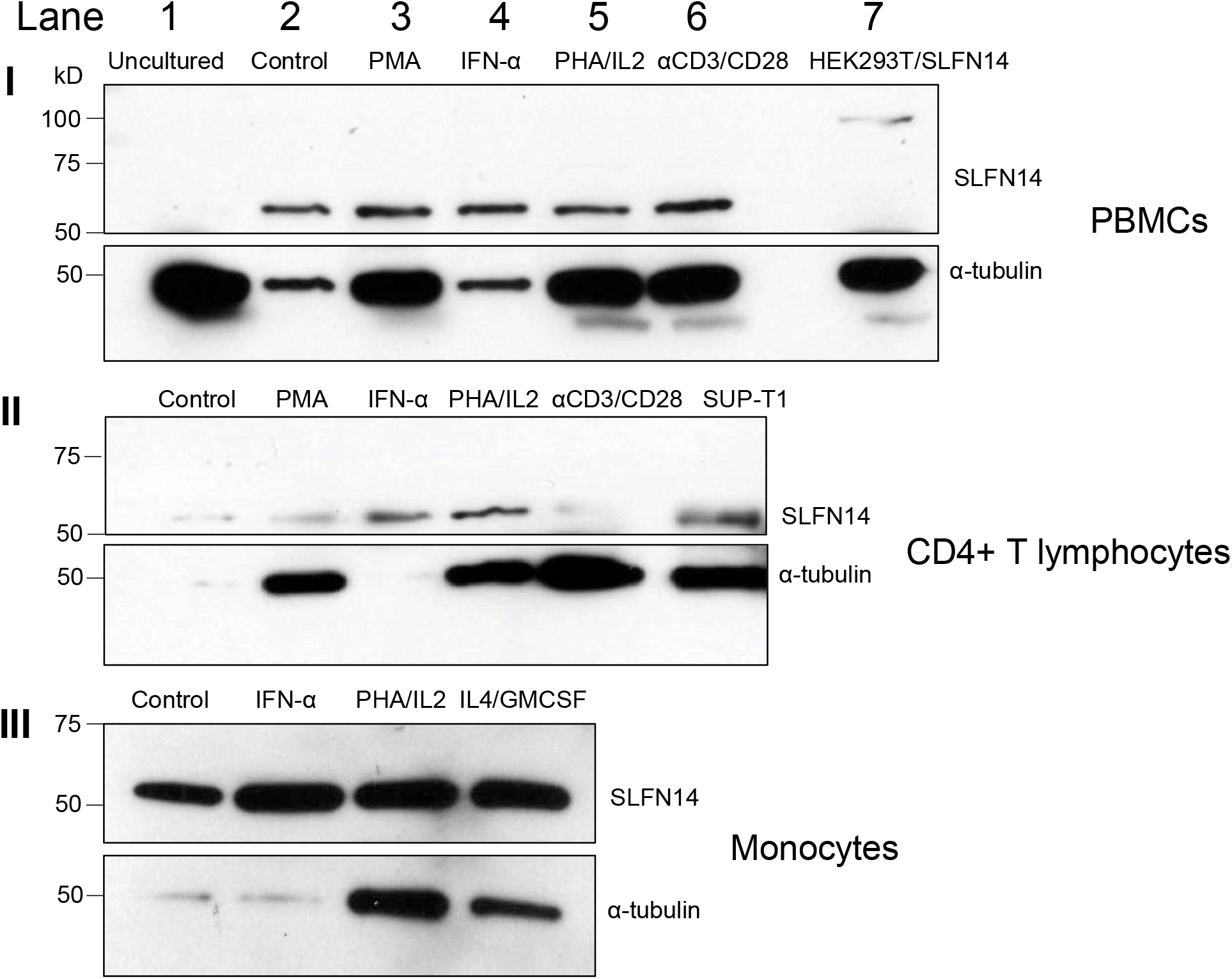
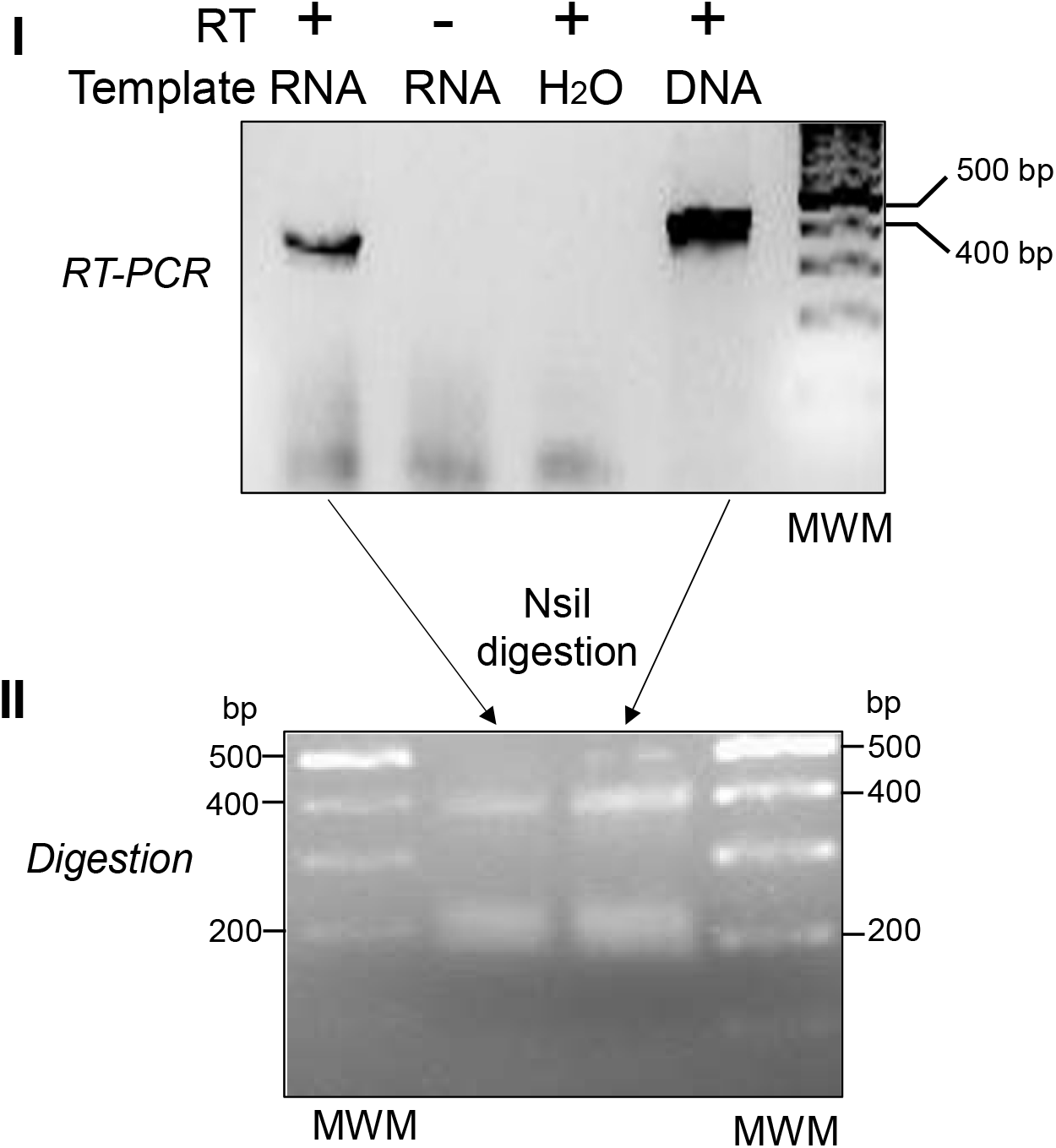
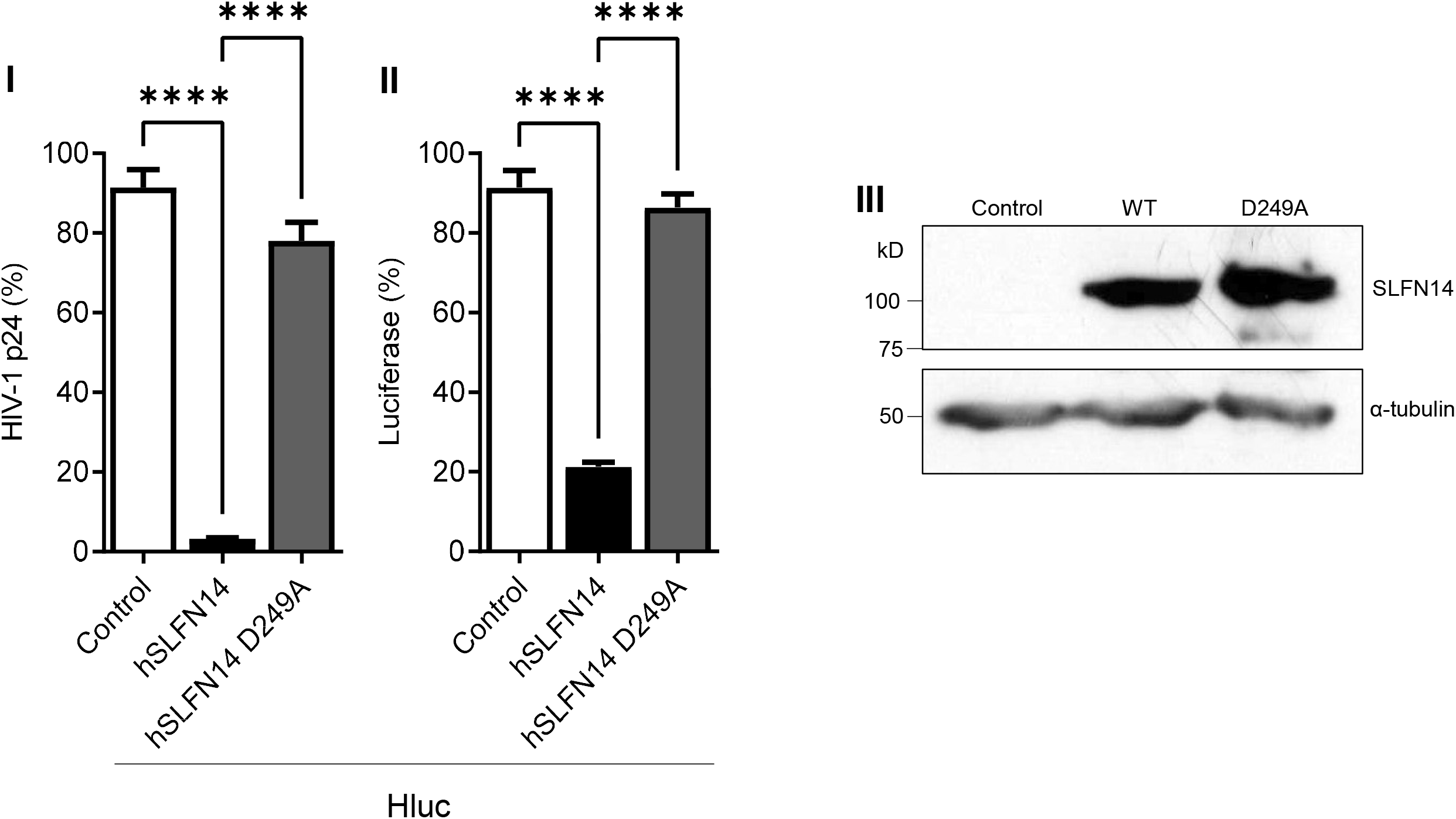
Expression of SLFN14 in immune cells. **(A)**. Immunoblot analysis of SLFN14 expression in HEK293T co-transfected with an empty plasmid (Control) or a plasmid expressing human SLFN14, and in non-transfected SUP-T1, CEM, and MOLT-3 cells. The MOLT-3 cell lysate was in the same immunoblot membrane than the other samples but separated by irrelevant samples that were removed from the image presented. Anti-SLFN14 antibodies recognizing the N- or the C-terminus of the protein were used as indicated, and α-tubulin levels were determined as a loading control with a specific antibody. **(B)** Immunoblot analysis of SLFN14 expression in PBMCs, and PBMC-derived CD4+ T lymphocytes and monocytes with an anti-C-terminal SLFN14 antibody. α-tubulin was determined as a loading control. Cells were subjected to the stimuli indicated. Positive controls were HEK293T cells co-transfected with human SLFN14 (HEK393T/SLFN14) and SUP-T1 cells. **(C)** Analysis of SLFN14 mRNA in MOLT-3 cells. **(I)**. Agarose gel electrophoresis analysis of the reverse transcription (RT)-PCR evaluating SLFN14 expression in MOLT-3 cells with primers SS1 and CV43. Samples were loaded in the gel in the following order: RT-PCR with RNA, No RT control (PCR with RNA), No template control (RT-PCR with no RNA), Positive control (RT-PCR with SLFN14 expression plasmid), and DNA Molecular Weight Marker (Thermo Fisher, FERSM1331). **(II)**. Agarose gel electrophoresis analysis of the digestion with NsiI of the RT-PCR products obtained with primers CV43 and SS1 using as template the SLFN14 expression plasmid and RNA extracted from MOLT-3 cells. DNA molecular weight markers are indicated. **(D)** Effect of D249A mutation on the activity of human (h) SLFN14 on transgene expression. HEK293T cells that were co-transfected with a plasmid encoding an HIV-1 virus expressing firefly luciferase and either an empty plasmid (control cells) or a plasmid expressing human SLFN14 WT or D249A mutant. HIV-1 p24 levels **(I)** and luciferase activity **(II)** were expressed as % of control cells. **(III)** SLFN14 expression was validated by immunoblot as described in Fig. 1A-III. Statistically significant differences were calculated with two-way ANOVA Dunnett *post hoc* tests. **** *P* ≤ 0.0001. Data correspond to a triplicate experiment and are representative of three independent experiments.

In addition, SUP-T1 cells (lymphoblastic lymphoma-derived CD4+T cell line), and the acute lymphoblastic leukemia-derived CD4+T cell lines CEM and MOLT-3 were evaluated by immunoblot with these antibodies. Data in Fig. 3A indicate that the anti-C-terminal SLFN14 antibody (I), but not the antibody directed against the N-terminus (II), reacted with a band that migrated slightly above 50 kD in the three cell lines, suggesting the expression of an N-terminal truncated form of the protein.

We also analyzed the expression of SLFN14 in primary immune cells. PBMCs were isolated by sedimentation on Ficoll-Paque, lysed, and proteins analyzed by immunoblot with anti-SLFN14 antibodies recognizing the C- and the N-terminus. No SLFN14 protein was detected in freshly isolated, uncultured PBMCs (Fig. 3B-I lane 1).

Furthermore, we explored the effect of different stimuli on the expression of SLFN14 in primary cells. Equal number of PBMCs (Fig. 3B-I), CD4+ T cells (Fig. 3B-II), or monocytes (Fig. 3B-III) were treated with culture medium alone (Control, Figs. 3B-I lane 2 and 3B-II and -III lane 1) or supplemented with phorbol myristate acetate (PMA, Figs. 3B-I lane 4 and 3B-II lane 2), interferon α1 (IFN-α1, Figs. 3B-I lane 4, 3B-II lane 3, and 3B-III lane 2), phytohemagglutinin and interleukin-2 (PHA/IL2 Figs. 3B-I lane 5, 3B-II lane 4, and 3B-III lane 3), anti-CD3/-CD28 antibody-coated beads (αCD3/CD28, Figs. 3B-I lane 6 and 3B-II lane 5), or interleukin-4 and granulocyte-macrophage colony-stimulating factor (IL4/GMCSF, Fig. 3B-III lane 4).

After three days of treatment all the cells in the culture were lysed in the same volume and equal volumes of the cell lysates were evaluated by immunoblot. Notice that rather than equal protein amounts, same number of initial cells in the culture were loaded per electrophoresis lane in these immunoblots. This strategy, combined with the detection of α-tubulin by immunoblot, additionally allowed verifying the effect of the treatments on cellular proliferation. As positive controls in these immunoblots were analyzed HEK293T cells transfected with SLFN14 (Fig. 3B-I lane 7) and SUP-T1 cells (Fig. 3B-II lane 6).

In contrast to fresh PBMCs (Fig. 3B-I lane 1), after three days of *in vitro* culture, a band of similar size that the one detected in CD4+ T cell lines (Fig. 3A and Fig. 3B-II lane 6) was observed in PBMCs (Fig. 3B-I lane 2) as well as in CD4+ lymphocytes (Fig. 3B-II lane 1) and monocytes (Fig. 3B-III lane 1). This band was detected only with the anti-C terminal SLFN14 antibody whereas the anti-N-terminal failed to recognize any protein in any of the primary cell lysates (Data not shown).

PMA, PHA/IL-2, and anti-CD3/-CD28 immunobeads induced robust cellular proliferation in PBMCs (Fig. 3B-I) and CD4+ T cells (Fig. 3B-II), as indicated by the α-tubulin levels detected in these cells, compared to the untreated cells (control). Similarly, PHA/IL-2, and IL-4/GMCSF induced cellular proliferation in monocytes (Fig. 3B-III). Considering α-tubulin levels as a proxy of the number of cells analyzed, stimuli that induced cell proliferation also decreased the expression of the truncated SLFN14 per cell (Fig. 3B). In contrast, IFN-α1 increased the levels of low molecular band reactive with the anti-SLFN14 antibody in CD4+ T cells (Fig. 3B-II) but not in PBMCs or monocytes (Fig. 3B-I and III).

To analyze a potential mechanism implicated in the lack of expression of full-length SLFN14 in MOLT-3 cells, we determined whether these cells express an SLFN14 mRNAs carrying nucleotides 386 – 788 of exon 3 that encode amino acids 115 – 248. This protein region contains the epitope recognized by the anti-N-terminal SLFN14 antibody that seems to be missing in the shorter protein recognized only by the anti-C-terminal SLFN14 antibody in immune cells. RT-PCR analysis (Fig. 3C-I) indicated robust expression of mRNAs harboring this region of exon 3, and as expected, no amplification was observed in the minus RT control, excluding inadvertent detection of genomic DNA. The identity of the RT-PCR product was determined by overlapping DNA sequencing (GenBank accession numbers OP548624 and OP548623), and by restriction digestion with NsiI (Fig. 3C-II). This enzyme is predicted to split the 402 bp RT-PCR product in 215 and 187 bp bands. In this experiment, partial digestion was obtained as indicated by the presence of the 402 bp RT-PCR product and a thick ∼200 bp band that we interpreted as the 215 and 187 bp bands closely migrating. Similar results were obtained with another set of primers targeting nucleotides 302 – 788 that encodes amino acids 86 – 248 in SLFN14 (Data not shown). These findings are in agreement with the reported exon/intron organization of SLFN14 (NM_001129820.2), excluding alternative splicing as a mechanism in the generation of the N-terminal deleted form of SLFN14 detected in immune cells. Therefore, a post-splicing event seems to determine the absence of SLFN14 full-length in these cells.

### SLFN14 requires the endoribonuclease activity to impair HIV-1 expression

Protein molecular weight prediction (Expasy) indicated that SLFN14 full-length protein is 104 kD, as evidenced by our immunoblots (for example Fig. 3A-I). A SLFN14 protein lacking the first 330 or 420 amino acids is estimated to be 69 or 56 kD, respectively, being in the molecular weight range of the N-terminal truncated form of SLFN14 detected in immune cells (Fig. 3A and 3B). These predicted N-terminal truncated SLFN14 proteins will lack the epitope recognized by the anti-N-terminal antibody (residues 100 – 250), including the residue 249 which is essential for the endoribonuclease activity [42].

Since cells expressing the N-terminal truncated form of SLFN14 are susceptible to HIV-1, we predicted that SLFN14 requires the endoribonuclease activity to repress expression of HIV-1 Gag. Therefore, we determined the anti-HIV-1 activity of an SLFN14 endoribonuclease-dead mutant (D249A) [42]. HEK293T cells were co-transfected with a plasmid expressing an HIV-1 reporter that expresses LTR-driven luciferase [54] and either the empty plasmid or plasmids expressing human SLFN14 wild type or the D249A mutant. In concordance with our previous observations, SLFN14 wild type decreases HIV-1 p24 (Fig. 3D-I) and luciferase (Fig. 3D-II) levels by approximately 98% and 80%, respectively. However, this inhibitory activity was drastically reduced by the D249A mutation, despite the wild type and mutant proteins were expressed at similar levels (Fig. 3D-III). Therefore, the endoribonuclease activity of SLFN14 is required for the role of this protein in gene expression regulation, and likely the N-truncated form of the protein expressed in HIV-1-permissive immune cells is inactive.

### SLFN14 impairs HIV-1 protein expression in CD4+ T cells and monocytes

To evaluate the effect of SLFN14 on HIV-1 protein expression in HIV-1-permissive immune cells, SUP-T1 cells and primary CD4+ lymphocytes and monocytes were electroporated with an empty plasmid (control) or a human SLFN14 expression plasmid, and a plasmid encoding wild type HIV-1 (pNL4-3). Three independent cultures of SUP-T1 cells and one culture of CD4+ lymphocytes per donor (two donors) were electroporated. Because of their low yield, monocytes from the two donors were mixed right before electroporation and electroporated as one culture. Seventy-two hours after electroporation, ATP levels were measured in triplicate in each of the cultures, and cell supernatant was transferred in triplicate to fresh, non-electroporated, SUP-T1 cells. Viral replication in the target SUP-T1 cells was evaluated by measuring HIV-1 p24 in the cell supernatant at different days post-transfer.

We found similar ATP levels in the cells electroporated with the empty or the SLFN14 expression plasmids (Fig. 4A), indicating similar cell viability in the HIV-1 producer cells. In contrast, viral production was severely impaired by SLFN14 in all the cells electroporated, as demonstrated by the differences in the HIV-1 replication curves observed in the SUP-T1 cells infected with the produced viruses (Fig. 4B). At the earliest collection time after viral transfer, SUP-T1 cells infected with virus produced in SUP-T1 cells co-electroporated with the control plasmid showed approximately 30-fold more virus than the cells infected with the virus from SUP-T1 cells co-electroporated with SLFN14 plasmid (Fig. 4B-I). These differences were around 3- and 22-fold in CD4 cells from donor 1 and in the monocytes, respectively (Fig. 4B-II and IV). However, we did not see differences in HIV-1 levels in SUP-T1 cells infected with the virus produced by the CD4 cells from the second donor at this early time point (Fig. 4B-III). Nevertheless, at the second collection time point SUP-T1 cells infected with the virus from CD4 cells from donor 2 co-electroporated with the control plasmid showed approximately 500-folds more virus than SUP-T1 cells infected with virus from the SLFN14 co-electroporated CD4 cells (Fig. 4B-III). Similarly, all the cells studied showed important differences at the second collection indicating that SLFN14 impaired viral production in the electroporated cells (Fig. 4B). At the third collection time no differences were observed in SUP-T1 cells infected with the virus produced in electroporated SUP-T1 or CD4 cells from donor 1 (Fig. 4B-I and II) and viral cytopathic effect were marked, correlating with the high levels of HIV-1 p24 (1000 – 500 ng/ml) of these cultures. In contrast, differences in viral replication persisted very markedly in SUP-T1 cells infected with virus produced in CD4 cells from donor 2 (∼10^4^-folds, Fig. 4B-III) and in monocytes (∼10^3^-folds, Fig. 4B-IV). These SUP-T1 cells exhibited less viral cytopathic effect and their HIV-1 p24 levels were between 80 to 1 ng/ml. Therefore, SUP-T1 cultures showing no differences at the last collection time likely were exhausted by viral replication. In sum, findings in Fig. 4 demonstrated that SLFN14 can also inhibit HIV-1 expression in cell types that are relevant *in vivo* to this virus.

**Fig. 4.**
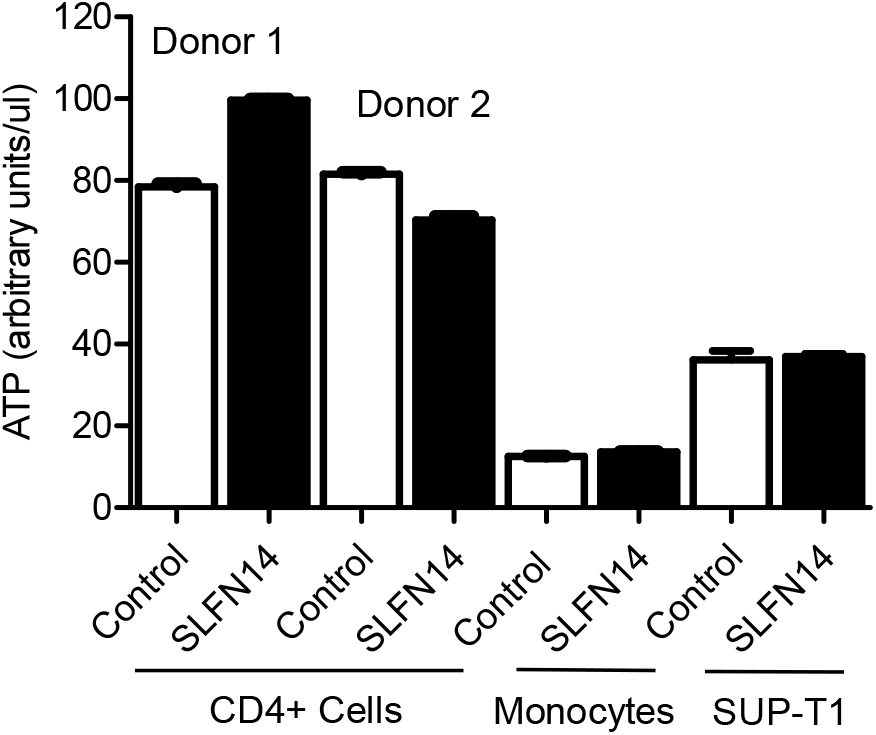
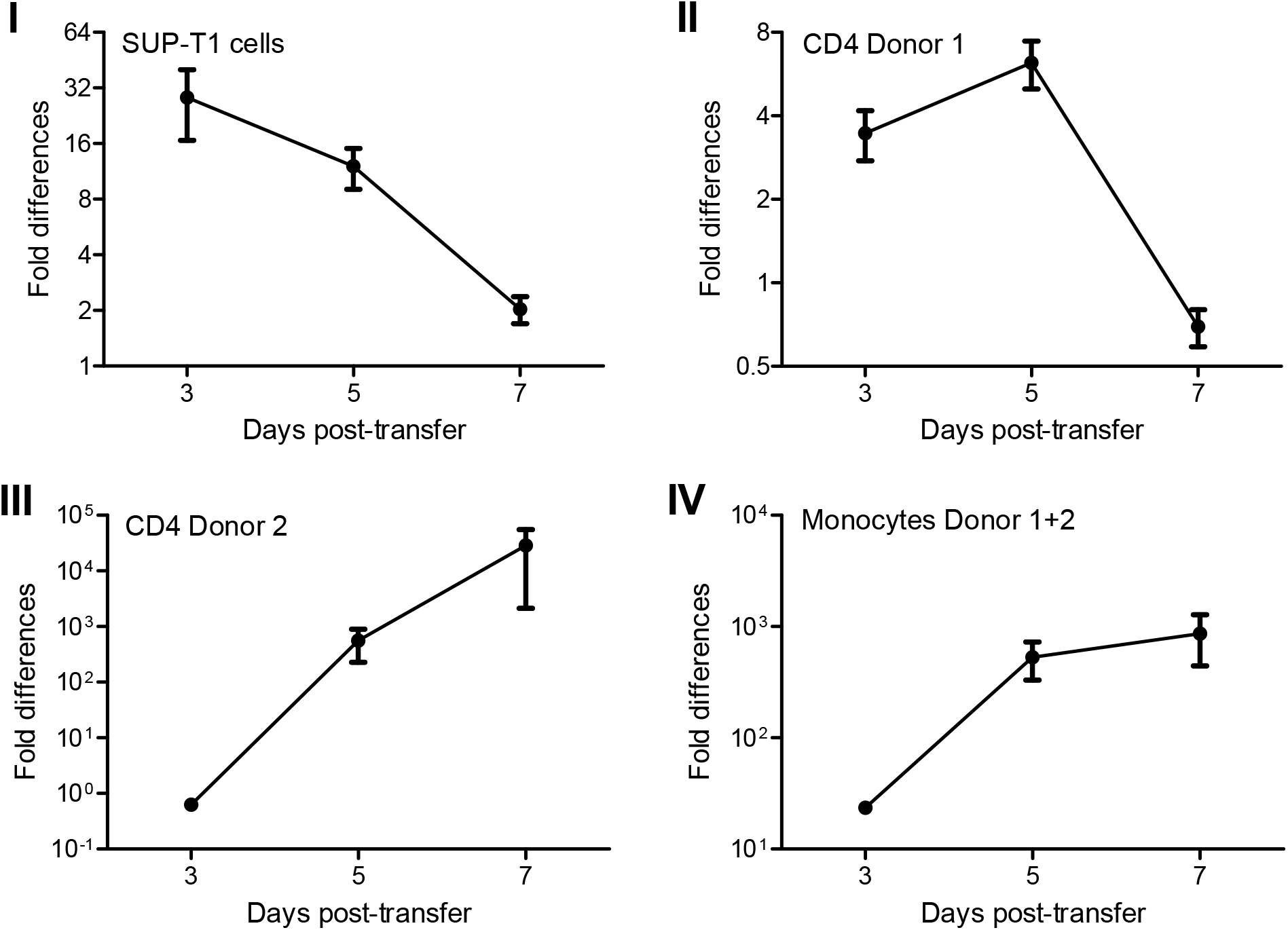
Effect of SLFN14 on HIV-1-driven protein expression in immune cells. Primary CD4+ T lymphocytes and monocytes, and SUP-T1 cells were electroporated with a plasmid expressing HIV-1 wild type and either an empty plasmid (control) or a plasmid expressing human SLFN14. Supernatant from the electroporated cells was transferred to fresh SUP-T1 cells and HIV-1 replication was followed by measuring HIV-1 p24 levels in the cell supernatant. **(A)** ATP levels in electroporated primary CD4+ T lymphocytes and monocytes, and SUP-T1 cells. **(B)**. Fold differences in HIV-1 p24 levels in SUP-T1 cells infected with the virus produced in electroporated SUP-T1 cells **(I)**, CD4+ lymphocytes from donor 1 **(II)** or donor 2 **(III)**, and donors 1 and 2 pooled monocytes **(IV)**. Data correspond to a triplicate experiment. Statistical analysis was conducted in by two-tailed, two-sample t test.

### Effect of SLFN14 on viral infection

Experiments reported above shown that SLFN14 impairs expression of rare codons-enriched transcripts expressed from transfected plasmids. Since SLFN14 affects preferentially the expression of wild type (AT-rich) rather than codon optimized (AT-poor) Gag, the RNA Pol-III-RIG-I pathway could be implicated. RNA Pol III transcribes AT-rich DNA templates producing short AU-rich RNA fragments that induce type I IFN via RIG-I [55], and this innate immune pathway is active in HEK293T cells [56]. Furthermore, transfection of HEK293T cells with *in vitro* transcribed HIV-1 wild type-encoded RNAs has been reported to trigger type I IFN, and this effect was inhibited by codon optimization of the viral sequences to resemble the human codon usage [57].

To investigate the implication of type I IFN signaling in the inhibitory effect of SLFN14 on gene expression, HEK293T cells were co-transfected with a plasmid encoding an HIV-1 reporter that expresses LTR-driven luciferase [54] and either, an empty plasmid or a plasmid expressing SLFN14. Transfected cells were treated with RNA polymerase III or pan-Janus kinase inhibitors during the entire duration of the experiment, and luciferase activity was measured in cell lysates 72 hrs after transfection. As expected, SLFN14 reduced by 80% the expression of luciferase found in control cells (Fig. 5A). This effect was not modified by RNA polymerase III or pan-Janus kinase inhibitors, indicating that the RNA Pol-III-RIG-I-IFN pathway is not involved in the ability of SLFN14 to impair gene expression.

**Figure 5.**
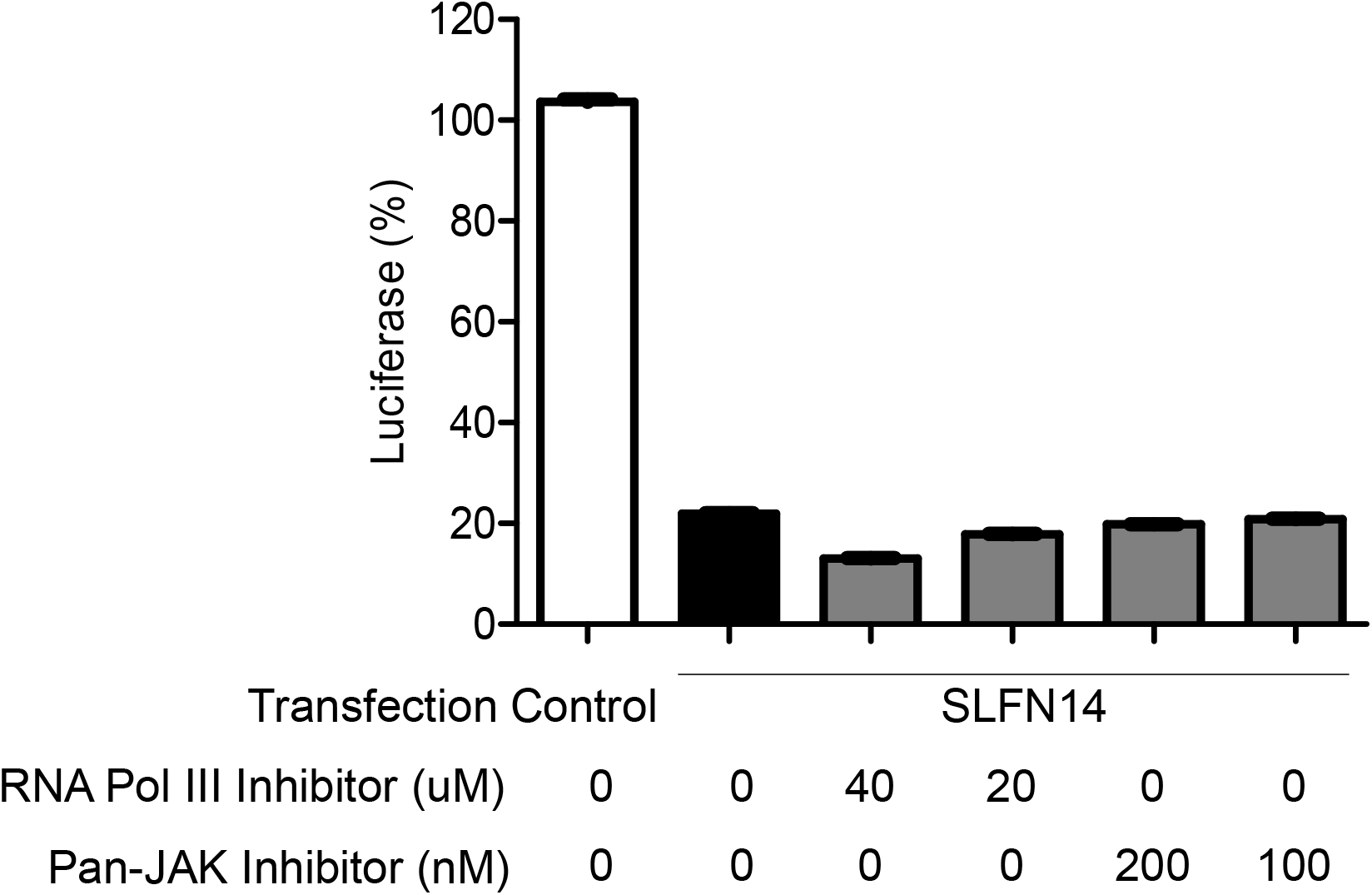
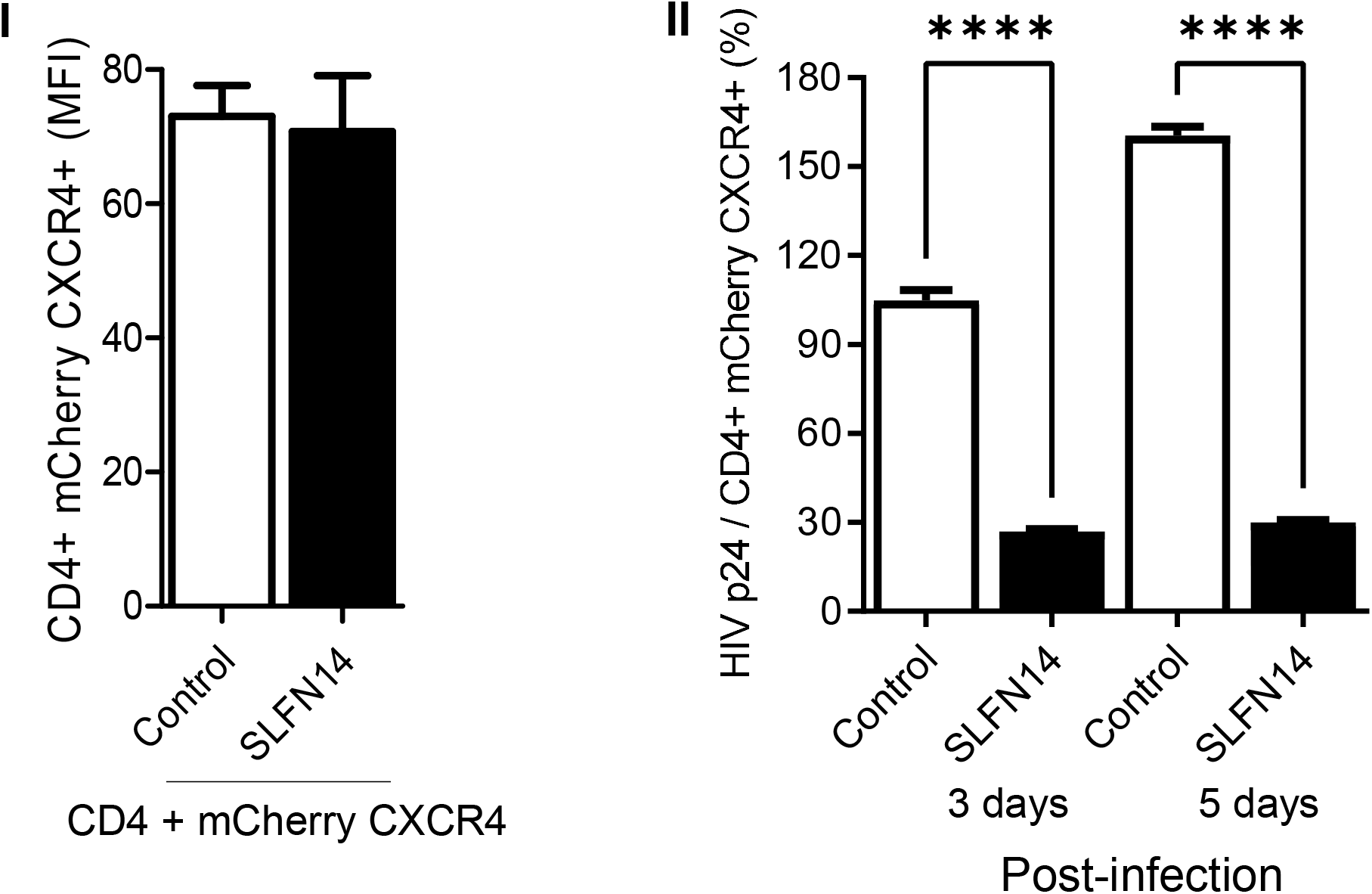
Effect of SLFN14 on viral replication. **(A)** Role of the RNA Polymerase III-RIG-I-IFN signaling pathway in the SLFN14 activity. HEK293T cells co-transfected with HIVluc expression plasmid and an empty plasmid (control cells) or a plasmid expressing human SLFN14 that were treated or not with inhibitors indicated. Luciferase activity was expressed as % of control cells in Data correspond to a triplicate experiment and are representative of three independent experiments. Although not indicated, statistically significant differences were found between the control and each of the other groups as calculated with two-way ANOVA Dunnett post-hoc test **** *P* ≤ 0.0001. **(B)** Effect of SLFN14 on HIV-1 replication. **(I)** HEK293T cells were co-transfected with plasmids expressing CD4 and a bicistronic plasmid encoding CXCR4 and mCherry, together with either an empty (control cells) or a human SLFN14 expression plasmid. MFI values of CD4 and mCherry (CXCR4). **(II)** These cells were infected with HIV-1 wild type and viral replication followed by measuring HIV-1 p24 in the cell supernatant. Data pertain to a triplicate experiment and they are representative of two independent experiments. Statistically significant differences were calculated with one-way ANOVA Bonferroni post-hoc test **** *P* ≤ 0.0001.

To evaluated the effect of SLFN14 on codon biased transcripts expressed independently of plasmid transfection, and since the inhibitory activity of SLFN14 is type I IFN-independent, we evaluated the effect of SLFN14 on the expression of rare codon-enriched transcripts expressed by viral infection. Then, we took advantage of HIV-1 again. Upon infection, HIV-1 inserts a cDNA copy of the viral genome in the host chromosome, and genes in this cDNA are expressed as any other gene in the cell. To maximize the frequency of HIV-1 infected cells containing the control or SLFN14 plasmids, we co-transfected HEK293T cells with plasmids expressing the HIV-1 receptor (CD4) and co-receptor (CXCR4) with either, plasmids empty or encoding SLFN14. Ectopic expression of HIV-1 receptor has been successfully used to study different aspects of HIV-1 biology [for example [58–60]]. In addition, we used this cellular model because the high transfection efficiency of HEK293T cells and their lack of endogenous CD4 and SLFN14 expression.

Forty-eight hrs after transfection, cells were infected with HIV-1 wild type (NL4-3 strain). Input virus was removed by extensive washing 24 hrs after infection, and cell supernatant collected at 0 (after wash), 3 and 5 days after infection to measure HIV-1 p24 by ELISA. Surface CD4 expression was measured at the time of infection by immunostaining with a specific antibody, and since CXCR4 was expressed from a bicistronic plasmid together with mCherry, levels of the fluorescence protein served as a surrogated of CXCR4 expression. HIV-1 p24 levels were normalized to the % of CD4+/mCherry CXCR4+ cells to account for transfection efficiency.

The MFI of CD4 and mCherry was similar in cells transfected with control and SLFN14 expression plasmids (Fig. 5B-I), indicating that SLFN14 did not affect the expression levels of CD4 or mCherry that exhibit CAIs of 0.82 and 0.976, respectively. However, cells expressing SLFN14 produced approximately ∼73% and ∼83% less HIV-1 p24 than control cells at days 3 and 5 post-infection, respectively (Fig. 5B-II). HIV-1 p24 at day zero was undetectable. These findings demonstrated that SLFN14 can impair expression of Gag from an integrated provirus.

### SLFN14 caused ribosomal RNA degradation in cells co-expressing Gag wild type

SLFN14 is a ribosome-associated endoribonuclease [41, 42]. Purified SLFN14 was reported to degrade purified ribosomal, tRNA, and mRNA *in vitro* [42]. In addition, our results here indicate that SLFN14 restricts the expression of transcripts rich in rare codons at a post-transcriptional step and requiring the endoribonuclease activity. Therefore, we sought to evaluated the effect of SLFN14 on rRNA and mRNA levels. In these studies, we used as a model wild type and codon optimized Gag, expecting to identify a SLFN14-dependent mechanism that operates preferentially in the cells expressing wild type Gag.

HEK293T cells were co-transfected with plasmids expressing HIV-1 Gag wild type or codon optimized together with a plasmid expressing firefly luciferase (pCI Luc), and either empty plasmid or plasmids expressing human or mouse SLFN14. Seventy-two hours after transfection, supernatant from these cells was used to measure HIV-1 p24, and cell lysates were evaluated for luciferase activity and the levels of SLFN14 proteins (immunoblot). Total RNA and DNA were also extracted from these cells to measure ribosomal RNA integrity by gel electrophoresis, Gag mRNA levels by quantitative reverse transcription PCR, and Gag DNA by quantitative PCR.

Similar amounts of mouse and human SLFN14 were detected by immunoblot in cells expressing wild type or codon optimized Gag (Fig. 6A). In contrast, SLFN14 proteins decreased wild type Gag expression by over 500 folds as compared to control cells, whereas codon optimized Gag expression was diminished only ∼2 folds (Fig. 6B). Luciferase levels were also diminished by SLFN14 proteins. In cells co-expressing Gag wild type, luciferase dropped by 17 folds and in cells co-expressing Gag codon optimized, luciferase was diminished by ∼2 folds (Fig. 6B). The fact that the expression of the same luciferase transcript was ∼8 folds more affected in cells expressing non-optimized (wild type) than optimized Gag is intriguing. This could be due to differences in transfection efficiency, although this possibility seems to be unlikely because equal levels of SLFN14 were observed in cells expressing Gag wild type and codon optimized (Fig. 6A). Nevertheless, after normalization for the effect of SLFN14 proteins on luciferase, still SLFN14s impaired expression of Gag wild type by ∼36 folds but Gag codon optimized expression was not affected (0.9 folds).

**Figure 6.**
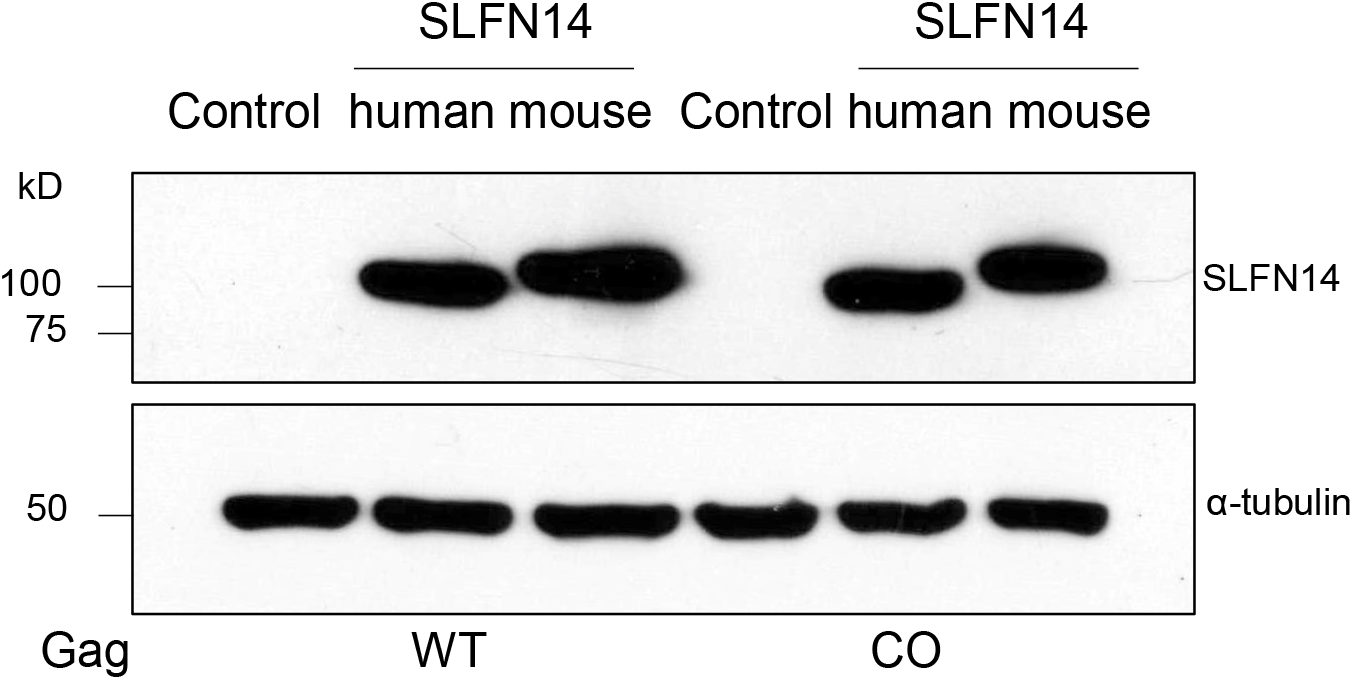
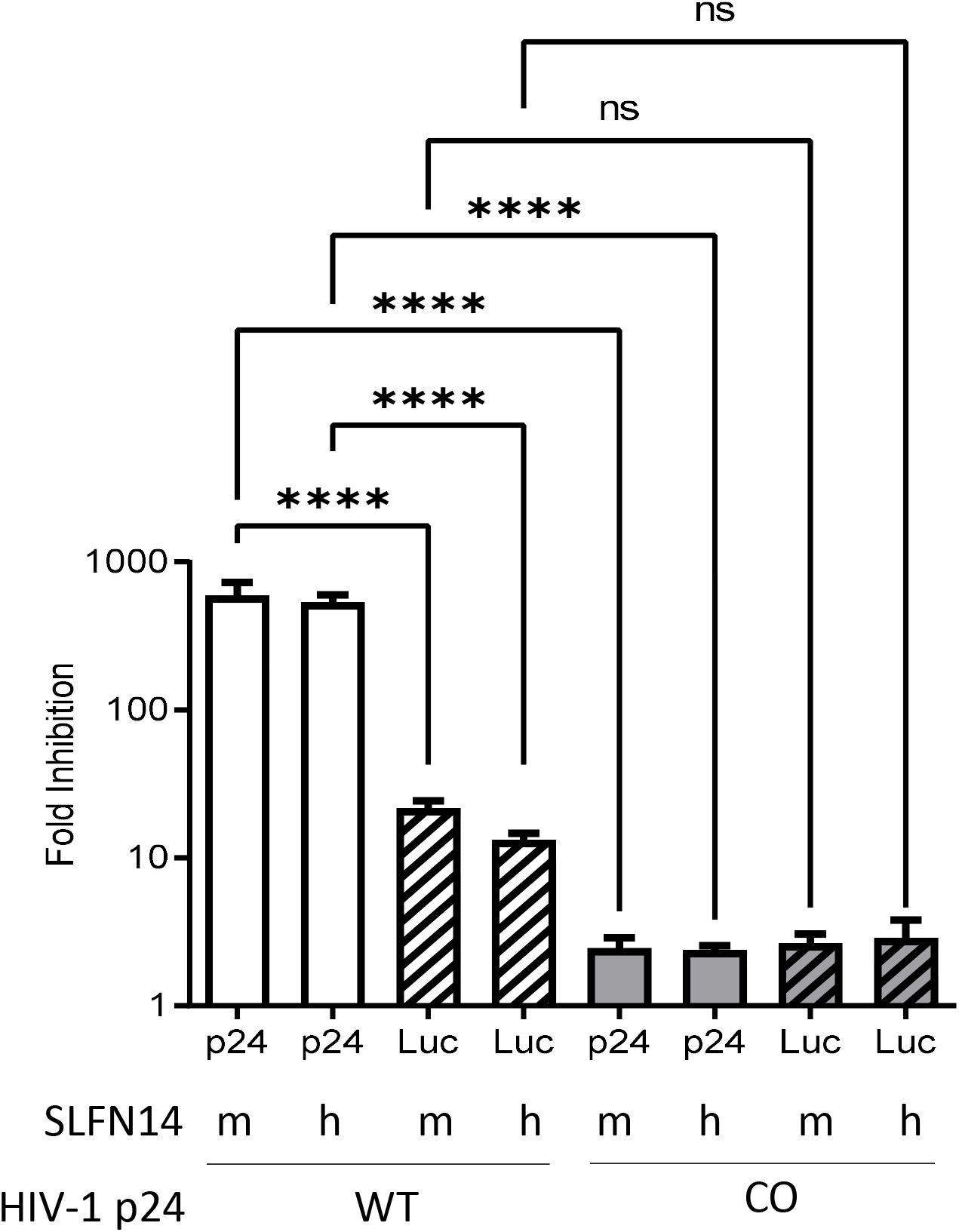
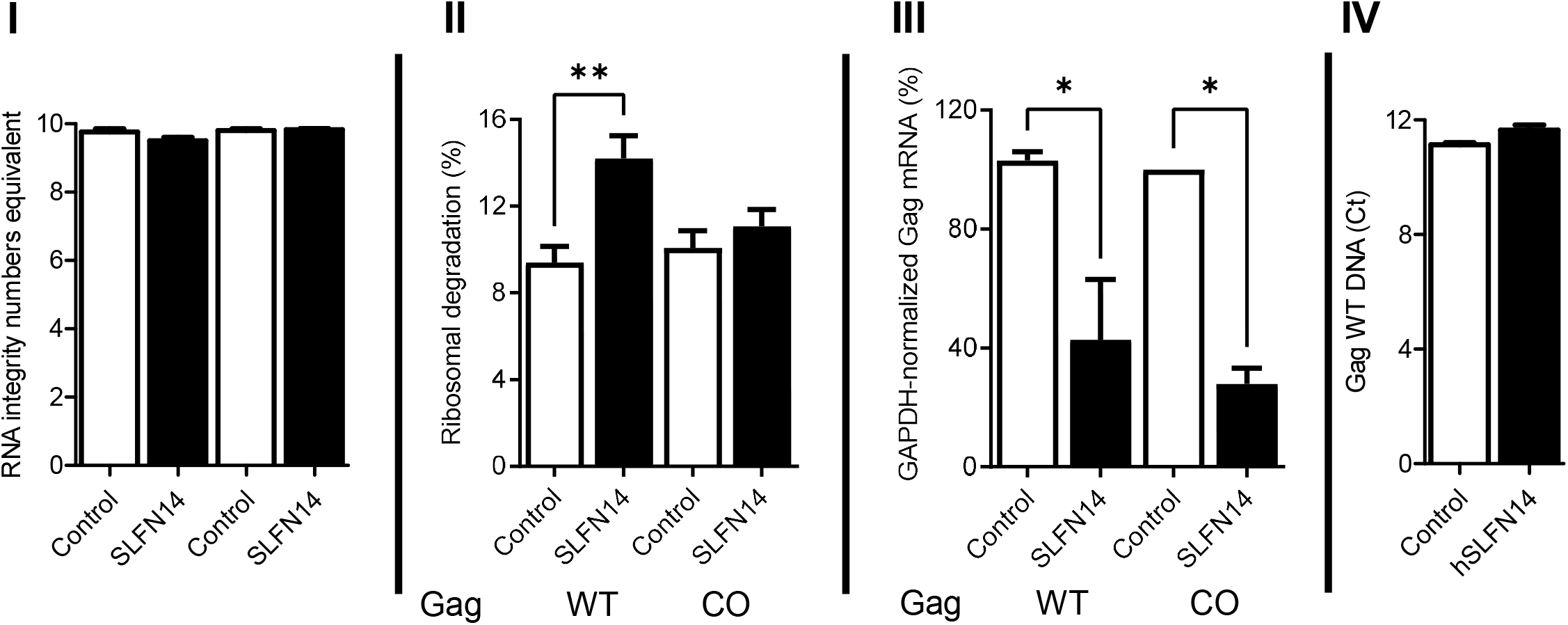
Effect of SLFN14 on nucleic acid integrity. HEK293T were co-transfected with a plasmid encoding Gag wild type (WT) or codon optimized (CO), and plasmids expressing luciferase (Luc) and either an empty plasmid (control cells) or a plasmid encoding SLFN14, human or mouse. **(A)** SLFN14 expression was verified by immunoblot as described in Fig. 1A-III. **(B)** HIV-1 p24 levels and luciferase activity determined in these cells were expressed as fold inhibition relative to control cells. Note that the Y axes of the graphic is in log_10_ scale. Data correspond to a triplicate experiment that is representative of five independent experiments. Statistically significant differences were calculated using one-Way ANOVA and a Tukey pos hoc test. **(C)** Effect of SLFN14 on nucleic acids. RNA quality **(I)**, ribosomal RNA degradation **(II)**, GAPDH mRNA-normalized Gag mRNA levels **(III)**, and Gag WT cDNA levels **(IV)** were expressed relative to values found in control cells. Data correspond to a triplicate experiment and is representative of two independent experiments. Statistically significant differences were calculated with one-way ANOVA and Dunnett *post hoc* tests. **** *P* ≤ 0.0001, *** *P* ≤ 0.001, ** *P* ≤ 0.01, * *P* ≤ 0.05, and NS *P* > 0.05.

After reverification of the differential effect of SLFN14 on the expression of wild type and codon optimized Gag, we proceeded to evaluate the effect of SLFN14 on the stability of nucleic acids obtained from these cells. Importantly, purified total RNA exhibited similar RNA Integrity Numbers equivalent (RIN^e^) (Fig. 6C-I) indicating a comparable high quality [61]. This total RNA was used to calculate ribosomal RNA degradation with the Agilent TapeStation Controller Software 4.1 following criteria previously reported [42, 62]. As compared to the corresponding control cells, cells transfected with SLFN14 showed a 1.7- or 1.1-fold increase in ribosome degradation when co-expressed with wild type or codon optimized Gag, respectively (Fig. 6C-II). These findings suggest that ribosome degradation could mediate the differential effects of SLFN14 on the expression of wild type and codon optimized Gag.

We also measured in these total RNA samples the levels of Gag mRNA using a set of primers that bind to the same region in Gag wild type and codon optimized, and values were normalized to GAPDH mRNA levels in the same samples. SLFN14 diminished by 2- and 3-fold the mRNA levels in cells co-expressing Gag wild type and codon optimized, respectively (Fig. 6C-III). Similar degradation of Gag mRNA in these cells cannot explain the differential effect of SLFN14 on the expression of Gag wild type and codon optimized, but suggest an additional mechanism for the inhibitory effect of SLFN14 on the expression of wild type and codon optimized HIV-1 p24 expression.

Because SLFN14 impaired the expression of genes located in plasmids or in the HIV-1 genome integrated in the host chromosome, unlikely, DNA degradation is implicated in the SLFN14 inhibitory activity. Nonetheless, levels of Gag wild type expression plasmid were determined in cells studied in Fig. 6 by quantitative PCR. Gag DNA cycle threshold (Ct) were 11.1 +/-0.1 in control cells and 11.6 +/-0.2 in SLFN14 cells (Fig. 6C-IV), indicating that the differences in HIV-1 p24 observed in these cells (Fig. 6B) were not due to different amounts of the Gag plasmid, excluding DNA degradation as a mechanism for these differences.

### Subcellular distribution of SLFN14

Since SLFN14 affect the expression of rare codon biased mRNAs at a post-transcriptional step and this is associated to ribosomal degradation, we also determined the subcellular distribution of human and mouse SLFN14 by immunostaining and confocal microscopy analysis. HEK293T cells were transfected with plasmids expressing mouse or human SLFN14 or an empty cassette, and a plasmid encoding a single-round infection HIV-1 expressing eGFP (HIVeGFP) [51]. Transfected cells were stained with an anti-FLAG antibody. As expected, SLFN14 proteins were distributed exclusively to the cell cytoplasm (Fig. 7A), and their localization did not change in cells co-expressing HIV-1 proteins (eGFP+ cells).

**Fig. 7.**
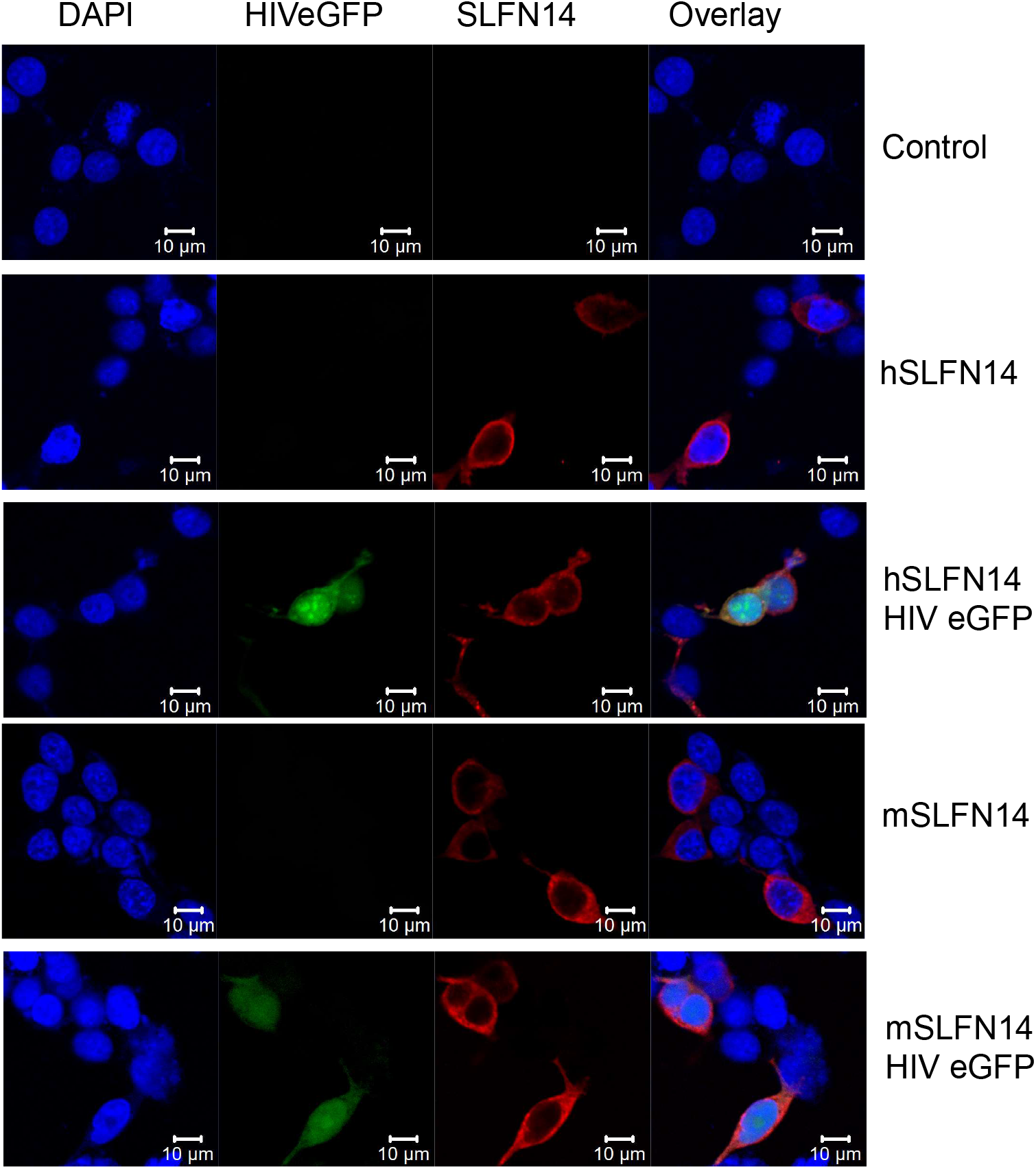
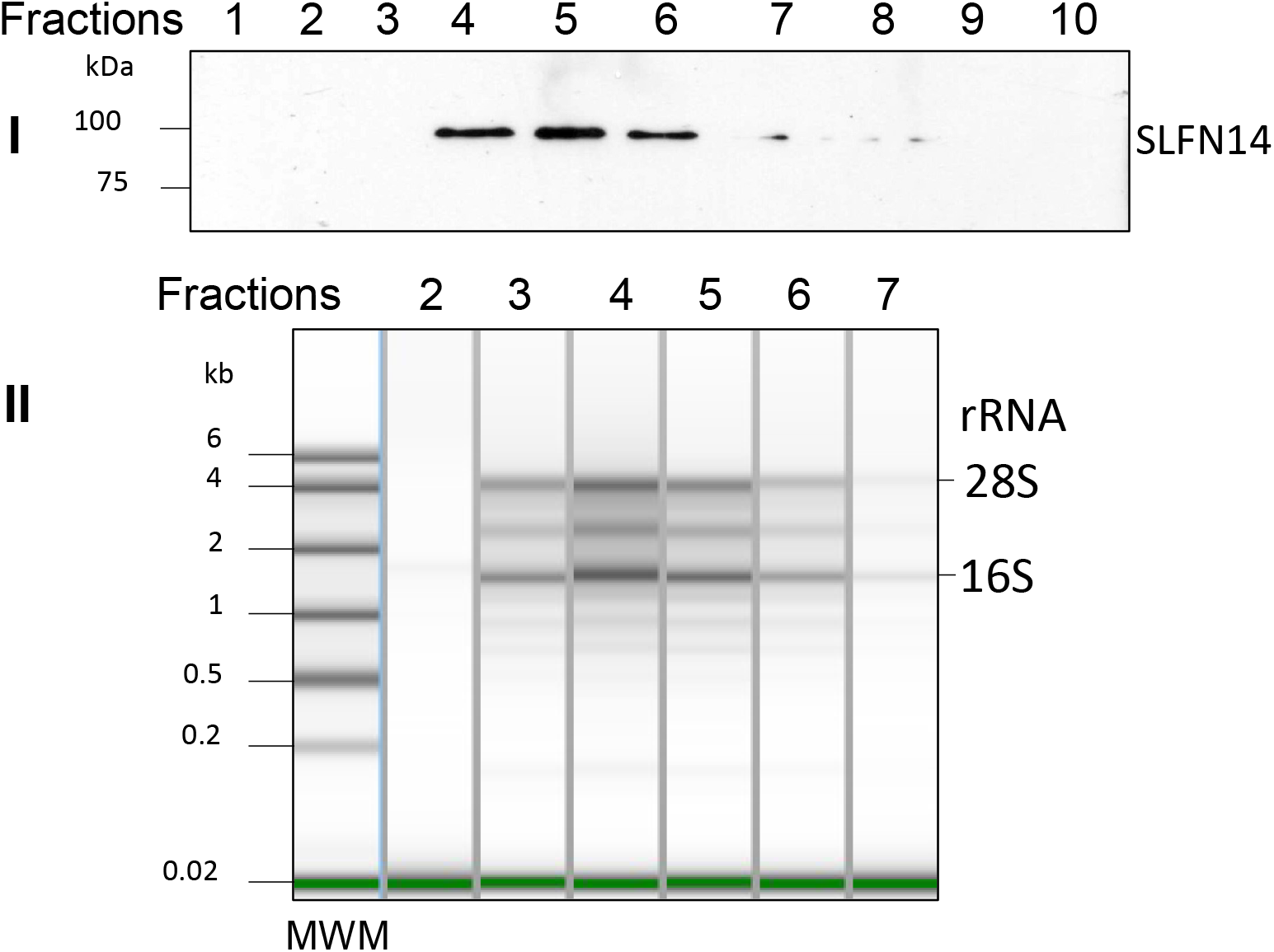
Ribosome association of SLFN14. **(A)** Subcellular distribution of SLFN14. SLFN14 was detected with an anti-FLAG antibody (red fluorescence) in HEK293T co-transfected with plasmids expressing mouse (m) and human (h) SLFN14 or an empty plasmid (control cells), and HIVeGFP expression plasmid or not. Nuclei were stained with DAPI. Data in the figure correspond to more than 100 cells in different fields of one experiment and are representative of three independent experiments. **(B)** Co-sedimentation of SLFN14 with ribosomes. **(I)** Immunoblotting detection of SLFN14 with an anti-FLAG antibody in sucrose gradient fractions obtained from HEK293T co-transfected with plasmids expressing human SLFN14 and HIV-1 Gag wild type. **(II)** The presence of ribosomes in selected fractions was determined by RNA gel electrophoresis. Data in the figure correspond to one experiment and are representative of five independent experiments

Furthermore, we verified the association of SLFN14 with ribosomes by sedimentation analysis in a sucrose density gradient, as previously described [41, 42]. HEK293T cells were co-transfected with plasmids expressing human SLFN14 or an empty plasmid and a plasmid expressing HIV-1 wild type Gag. The cells were lysed, and the cytoplasmic fraction was resolved in a sucrose density gradient. All the fractions obtained were analyzed for the presence of SLFN14 by immunoblotting with an anti-FLAG antibody but this protein was visible only in the middle region of the top quarter of the gradient (Fig. 7B-I). Gel electrophoresis analysis of RNA isolated from the SLFN14-containing fractions and the corresponding flanking fractions lacking the protein indicated the presence of SLFN14 in ribosomal RNA-enriched fractions (Fig. 7B). Therefore, SLFN14 co-sedimented with ribosomal fractions, in concordance with previous reports [41, 42].

## DISCUSSION

The SLFN family is widely distributed in mammals [63, 64]. In mouse and humans, different genes compose this family and they are classified according to their protein length in three groups. Human SLFN group III proteins are implicated in translational control, and these functions mediate their antiviral activities [32, 39, 65]. SLFN11, and SLFN13 bind and degrade tRNAs [38, 39, 66], whereas SLFN5 binds tRNA without causing their degradation [40], and, *in vitro*, SLFN14, a ribosome-associated protein, degrades rRNA, tRNA, and mRNA [42]. Similarly, to SLFN5, the group II protein SLFN2 binds tRNA protecting them from stress-induced, angiogenin-mediated degradation [28]. Our findings further expand the role of this family in translational control by demonstrating in cells for the first time that SLFN14 impairs expression of transcripts enriched in rare codons at a post-transcriptional step, likely at translation. SLFN14 drastically reduced the protein levels of wild type HIV-1 Gag and firefly luciferase that exhibit a low CAI of 0.56 and 0.71, respectively, but did not affect the expression of proteins encoded by transcripts enriched in codons frequently used in human cells, such as codon optimized HIV-1 Gag (CAI 0.99), HIV-1 Tat (CAI 0.761), CFP (CAI 0.96), mCherry (CAI 0.976), eGFP (CAI 0.962), and CD4 (CAI 0.82). Furthermore, SLFN14 more drastically impaired expression of wild type Gag than luciferase, correlating with the lower CAI of Gag. SLFN14 was also reported to impair the expression of other viral proteins such as the varicella zoster virus proteins immediately early 62 (CAI 0.72) and glycoprotein E (CAI 0.7) [44, 45], and the nucleoprotein from influenza virus (CAI 0.753) [45]; in support of our hypothesis these viral proteins also have a low CAI. Therefore, we propose that SLFN14 regulates gene expression in a rare codon-dependent manner.

Importantly, the effect of SLFN14 on HIV-1 protein expression was observed in several cell types, including primary cells implicated in HIV-1 infection *in vivo* (CD4+ T cells and monocytes) and in SUP-T1 and HEK293T cells; and on HIV-1 transcripts generated by plasmid transfection or by viral infection. Because the base composition of the HIV-1 genome is very stable over time, varying less than 1% per base per isolate regardless the HIV-1 group of subtype or the time of isolation during the pandemic [57], SLFN14 could be a restriction factor for HIV-1 infection.

Mechanistically, the anti-HIV-1 activity of SLFN14 requires the endoribonuclease activity of this protein and when co-expressed with HIV-1 Gag wild type, but not with a codon optimized version of this viral protein, SLFN14 triggered ribosome degradation. In contrast, Gag mRNA was similarly degraded in cells co-expressing SLFN14 and Gag wild type or codon-optimized, indicating that mRNA degradation cannot explain the more potent inhibitory effect of SLFN14 on wild type than in codon optimized Gag expression. Then, it is possible that, in response to translation elongation constraints experienced during the translation of codon biased mRNAs, the ribosomal-resident protein SLFN14 triggers degradation of ribosomes implicated in the translation of these messengers.

SLFN14 could also act synergistically with SLFN11, and potentially SLFN13, to modulate the expression of transcripts with bias towards rare codons. In this functional crosstalk, SLFN11 and/or SLFN13 would decrease tRNA abundance globally, increasing the translation elongation constraints experience by rare codon-enriched transcripts, making these mRNAs more susceptible to SLFN14 restriction.

Diverse cellular proteins are encoded by transcripts exhibiting codon usage bias [67]; therefore, if SLFN14 represses the expression of transcripts of this type, tight control of the SLFN14 activity or expression seems to be important. Our data indicate that SLFN14 protein activity is regulated by limiting the expression of the protein (i.e., HEK293T and uncultured PBMCs) or by producing a N-terminal truncated form, as observed in cultured primary PBMCs, CD4+ T lymphocytes and monocytes, as well as in immortal CD4+ T cells. This N-terminal truncated form is predicted to lack a region of the protein that contains the catalytic residue D249; therefore, to be nonfunctional. The truncation mechanism is unclear, RT-PCR analysis of MOLT-3 cells, that express the truncated SLFN14, indicated the presence of the first nucleotides of the exon 3, that encodes the potentially missing residues. In addition, the GeneBank-reported exon/intron organization does not support an alternative splicing mechanism to generate the shorter SLFN14. Unfortunately, no anti-SLFN14 antibodies, other than the antibodies we used in this research, are commercially available to further verify the hypothesis of the N-terminus truncation.

In summary, our findings indicate a novel role of the ribosome-associated endoribonuclease SLFN14 in regulating transcript expression in a rare codon-dependent manner.

## MATERIALS AND METHODS

### Cell lines

SUP-T1, CEM and MOLT-3 cells and primary immune cells were grown in RPMI 1640 medium supplemented with 20% heat-inactivated fetal calf serum, 2 mM L-glutamine, and 1% penicillin-streptomycin, while HEK293T cells were grown in Dulbecco’s modified Eagle’s medium (DMEM) supplemented with 10% heat-inactivated fetal calf serum, 2 mM L-glutamine, and 1% penicillin-streptomycin. All the cell lines used were previously obtained from ATCC.

### Expression plasmids

pHluc was derived from pNL4-3luc-R-E- as described in [54]. This single-round infection HIV-1 expresses LTR-driven firefly luciferase from the *nef* slot, lacks VPR expression, and has a 426 nt deletion in the *env* gene. FLAG-tagged human and mouse SLFN14 (Origene, RC226257 and MR225976) were expressed from pCMV6-Entry. Empty plasmid was derived from the human SLFN14 expression plasmid by deleting the entire open reading frame (2.8 kb) by digestion Sal I / Xho I and religation of the backbone (4.8 kbs). FLAG-tagged human SLFN14 D249A was generated by site directed mutagenesis with the QuickChange Lightning Site-Directed Mutagenesis Kit (Agilent) using reverse primer SS1 (5’-atccaccccaatgaggacatatcc-3’) and forward primer SS2 (5’-gCtaagagcaaagaagtggttggatg-3’), the point mutation is indicated in upper case in the primer sequence. The entire sequence of the SLFN14 D249A cDNA was verified by overlapping DNA sequencing. Wild type Gag was expressed from pCMVΔR8.91 (a gift of D. Trono) and codon optimized Gag from pARP-8675 (NIH AIDS Reagent Program). This construct expresses a codon-optimized Gag pre-protein from HIV-1 clone 96ZM651.8 [68]. Cyan fluorescent protein (CFP) expression plasmid was pECFP-C1 (Clontech). Plasmids pCAGGS-CD4-Myc (Addgene, 58537), and pRP-mCherry/Puro-CAG>hCXCR4 (VectorBuilder, VB900125-2200scz) were used to express CD4 and CXCR4 respectively. The CXCR4 expression plasmid contains an independent mCherry expression cassette (bicistronic plasmid). pCI Luc contains firefly luciferase cDNA cloned MluI / Xba I in pCI (Promega). pNLENG1-ES-IRES (a gift of D.N. Levy, NYU) encodes a single-round infection HIV-1 (HIVeGFP) that contains two stop codons in *env* and the eGFP open reading frame is inserted between the *env* and *nef* sequences. [69]. pNL4-3 encodes a wild type HIV-1 (strain NY5/BRU, LAV-1).

### Generation of viruses

Procedures previously described [70, 71] were used for production of NL4-3 HIV-1 virus. Briefly, HEK293T cells were co-transfected by calcium-phosphate with 15 ug of pNL4-3. Seventy-two hours after transfection the viral supernatant was harvested.

### Immunoblotting

HEK293T cells (∼3×10^6^) were lysed in 100 ul of Laemmli sample buffer (12 mM Tris-Cl, pH 6.8, 0.4% SDS, 2% glycerol, 1% β-mercaptoethanol, 0.002% bromophenol blue). SUP-T1, CEM and MOLT-3 cells, peripheral blood mononuclear cells (PBMCs), CD4+ T lymphocytes, and monocytes were lysed for 15 min on ice in 100 μl of CSK I buffer [72] (10 mM PIPES [piperazine-*N*,*N*′-bis(2-ethanesulfonic acid)] (pH 6.8), 100 mM NaCl, 1 mM EDTA, 300 mM sucrose, 1 mM MgCl_2_, 1 mM dithiothreitol (DTT), 0.5% Triton X-100) containing protease inhibitors (final concentrations of 2 μg/ml leupeptin, 5 μg/ml aprotinin, 1 mM phenylmethylsulfonyl fluoride, and 1 μg/ml pepstatin A). Cell lysates were centrifuged at 22,000 × *g* for 3 min at 4°C, and the supernatant mixed with Laemmli sample buffer, boiled for 10 mins, and saved at -80LJC for further analysis. Cell lysates (15 ul) was resolved by SDS-PAGE and transferred overnight to polyvinylidene difluoride (PVDF) membranes at 100 mA at 4°C. Membranes were blocked with Tris-buffered saline (TBS) containing 10% milk for 1 h and then incubated with the corresponding primary antibody diluted in TBS-5% milk-0.05% Tween 20 (antibody dilution buffer). FLAG-tagged mouse and human SLFN14 was detected with anti-FLAG MAb (1/500) (M2; Sigma), non-tagged human SLFN14 was detected with antibodies anti-SLFN14 PAb (Abcam, ab254806) (1/500) and anti-SLFN14 PAb (Invitrogen, PA520868) (1/500), that recognize epitopes in the N-terminal and C-terminal regions, respectively. As a loading control, anti-α-tubulin MAb (clone B-5-1-2; Sigma) was used at a 1/4,000 dilution. Membranes were incubated overnight at 4°C with anti-FLAG and anti-SLFN14 antibodies, whereas anti-α-tubulin MAb was incubated for 30 mins at 25°C. Primary antibody-bound membranes were washed in TBS-0.1% Tween 20, and bound antibodies were detected with goat anti-mouse Ig-horseradish peroxidase (HRP) (Sigma, 1/2,000) or mouse anti-rabbit IgG-HRP (Santa Cruz Biotech, 1/4,000) diluted in antibody dilution buffer. These antibodies were incubated for 1 hr at 25°C. Unbound secondary antibodies were washed as described above, and bound antibodies were detected by chemiluminescence.

### Analysis of SLFN14 activity

HEK293T cells were plated at 0.45 × 10^6^ cells/well in a six-well plate, or at 10^6^ cells in a T25 flask and transfected by calcium-phosphate with the corresponding plasmids, and transfection medium was replaced with fresh culture medium 18 hrs later. In experiments evaluating the effect of SLFN14 on protein expression, cells were transfected in six-well plates. Each well was transfected with 1 ug of empty plasmid or 1ug of mouse or human SLFN14 and 1ug of the target plasmid (pNLENG1-ES-IRES, pHLuc, pCI Luc, pECFP-C1, pCMVΔR8.9, or pARP-8675), and cells were analyzed 72 hrs after transfection. In experiments evaluating the effect of SLFN14 on the HIV-1 infection, cells were plated in T25 flasks and transfected with 5 ug of empty plasmid or mouse or human SLFN14. Forty-eight hrs after transfection cells were detached mechanically and infected with HIV-1 (NL4-3) by spin-inoculation by resuspending cells (∼2×10^6^) in 500 ul of 37LJC-warmed culture medium in a 15 ml tube and centrifuged at 1,200g for 2 hrs at room temperature. Input HIV-1 wild type was removed the next day by extensive washing in culture medium and viral replication was determined by p24 ELISA at days 3 and 5 post-infection.

### Purification of primary cells

Blood samples were obtained from two deidentified healthy individuals in concordance with approved protocol from the Institutional Biosafety Committee of the University of Texas at El Paso and after informed consent was obtained from all subjects. All experiments were performed in accordance with relevant guidelines and regulations of our institution. Blood (60 mls) was centrifuged at 600 xg for 10 mins and plasma removed. Blood cells were mixed with 6 mls of PBS and 25 mls of this cell suspension was layered on 18 mls of Ficoll-Paque (Fisher Scientific, 17144003) and spun down for 35 mins at 400 x g at room temperature. The PBMC fraction was harvested, diluted three-fold in PBS, and collected by centrifugation at 600 xg for 10 mins. Uncultured fresh PBMCs (10^6^) were lysed in 50ul of CSKI, as described above, an analyzed by immunoblot. PBMCs (0.33×10^6^) were plated in round-bottom wells in 96 wells-plates in 100ul of culture medium and treated with different stimuli for 72 hrs. In addition, PBMCs (4X10^6^) were subjected to isolation of naïve CD4+ T cells (MiltenyiBiotec 130-094-131) or monocytes (MiltenyiBiotec 130-096-537) following the manufacturer instructions, and isolated cells were plated in round-bottom wells in 96 wells-plates in 100ul of culture medium and treated with different stimuli for 72 hrs. Treatments were culture medium (control), IL2 (30 U/ml) plus PHA (5 ug/ml), PMA (100 ng/ml), and IFN-α1 (10,000 U/ml). PBMCs and CD4+ T cells were also treated with Anti-Biotin MACSiBead (MiltenyiBiotec 130-091-441) loaded with anti-CD3 and anti-CD28 antibodies (anti-CD3/CD28 immunobeads) using one bead per two cells, and monocytes with GM-CSF (50 ng/ml) plus IL4 (50 ng/ml). Cells from one well of the 96-wells tissue culture plate were lysed in 50ul of CSKI, as described above, an 15ul of the cell lysate analyzed by immunoblot.

### Electroporation of immune cells

SUP-T1 cells and primary CD4+ T lymphocytes and monocytes were electroporated using Amaxa Cell Line Nucleofector Kit V (VCA-1003), and programs O-017 (T cells) or V-001 (monocytes) with 1ug pNL4-3 and 1ug empty plasmid or human SLFN14 expression plasmid. Seventy-two hrs after electroporation cell viability was determined by measuring ATP levels (Promega G9241) and 30 ul of cell supernatant was transferred to fresh SUP-T1 cells (10^5^ cells in 500 ul). HIV-1 p24 was determined by ELISA in the SUP-T1 cell culture supernatant at different times post-transfer.

### RNA polymerase III (RNA Pol III) and Janus kinase (JAK) inhibitors

HEK293T cells were transfected as described above with empty or human SLFN14 expression plasmids and a plasmid encoding Hluc in the presence of RNA Pol III inhibitor (Sigma ML-60218, 40 and 20 uM) or tofacitinib (Sigma PZ0017, 200 and 100 nM). Transfection medium was replaced with fresh culture medium 18 hrs later and inhibitors were added again. Cells were analyzed 72 hrs after transfection.

### HIV-1 p24 ELISA

HIV-1 p24 levels were determined by a sandwich ELISA according to the manufacturer’s instructions (ZeptoMetrix, 0801002). Briefly, cell culture supernatants were diluted appropriately and incubated on the ELISA antibody pre-coated wells overnight at 37°C. Unbound proteins were removed by washing the wells 6 times with 200 μl of washing buffer, and bound HIV-1 p24 was detected by incubating each well with 100 μl of the anti-HIV-1 p24-HRP secondary antibody for 1 h. Unbound antibodies were removed by washing as described above, and bound antibodies were detected by incubating each well with 100 μl of substrate buffer for 30 min at room temperature until the reaction was stopped by adding 100 μl of stop solution into each well. The absorbance of each well was determined at 450 nm using a microplate reader (Versa max microplate reader; Molecular Devices).

### Luciferase assay

Cells in suspension (100 ul, ∼3×10^5^) were mixed with 75ul of 0.1% Triton X-100 in PBS and 25 μl of substrate (Bright-Glow™ Luciferase Assay System, Promega), and 50ul aliquots of the mix were distributed in triplicate wells of a 96-wells white plate and analyzed in a microplate luminometer.

### CD4 and CXCR4 expression by flow cytometry analysis

CD4 and CXCR4 expression was detected in HEK293T cells (10^6^) transfected with 5 ug of empty plasmid or plasmids expressing human or mouse SLFN14 together 5 ug of plasmids expressing human CD4 and CXCR4. Transfected cells were harvested by mechanical dissociation and 10^5^ cells re-suspended in 100 uL of 1x PBS containing 1 uL of Alexa 488-labeled anti-human CD4 (eBioscience™, 53-0048-42) and incubated on ice for 5 min. CXCR4 and mCherry are expressed from the same bicistronic plasmid and therefore mCherry was used as a proxy of CXCR4 transfection efficiency. Cells were analyzed with a Gallios flow cytometer (Beckman Coulter). Fluorescence minus one controls were used to set up the flow cytometer. Data was analyzed with Kaluza Analysis software version 1.3.

### Quantitative RT-PCR and PCR, and rRNA integrity analyses

Total RNA was isolated from cells or sucrose density gradient fractions using TRIzol LS reagent (Invitrogen 10296010) according to the manufacturer’s instructions. All RNA samples had ratios of absorbance at 260/280 nm of 1.8 to 2.0, indicating that samples were contaminant-free and RNA integrity numbers equivalent (RIN^e^) >9.0. Purified RNA samples were stored at -80°C until use. Different mRNAs were detected by RT-PCR (BioRad, 1725151) using 1 ng of total RNA. HIV-1 Gag wild type mRNA was detected with primers DR22 (5’-agcaggaactactagtaccc-3’) and DR23 (5’-ttgtcttatgtccagaatgc-3’), while primers CV35 (5’-cgccggcaccacaagcaccc-3’) and CV36 (5’-ctgcttgatgtccaggatgc-3’) were used to detect Gag codon optimized, and primers EL9 (5’-acccctggccaaggtcatcc-3’) and EL10 (gacggcaggtcaggtccacc) were used for GAPDH. Human SLFN14 was detected by RT-PCR using primers CV43 (5’-atggatgttttcagccttccactaaggatttgc-3’) or SS3 (5’-GCTCCTTCCTTCAGGTTCACAG-3’) and SS1 (5’-atccaccccaatgaggacatatcc-3’) that bind to exon 3 which encodes the N-terminal region of the protein. DNA was extracted from cells transfected with pCMVΔR8.91 using TRIzol LS reagent and PCR amplified with primers DR22 and DR23 using IQ SYBR Green Supermix (BioRad, 1708882). Total RNA from 0.33×10^6^ cells was used for electrophoretic analysis of rRNA integrity with an Agilent Technologies 4200 TapeStation. As previously reported [62], RNA degradation bands were considered those migrating between the 18S rRNA and small RNAs bands in the region of 1500 base pairs (bp) to 200 bp. RNA degradation bands were quantified with the Agilent TapeStation Controller Software 4.1.

### Sucrose Density Gradient

HEK293T cells (3X10^6^) were plated in a T25 and calcium-phosphate transfected with 4 ug of empty plasmid or human SLFN14 expression plasmid and 4 ug of pCMVΔR8.91 or pARP-8675. The next day the transfection medium was replaced with fresh culture medium and 48 hrs later cells were mechanically harvested and used for the sucrose density gradient (SDG), as previously described [41, 42]. Briefly, cells were lysed by incubation in ice in Buffer A (20 mM Tris-HCl, pH 7.5, 100 mM KCl, 2.5 mM MgCl 2, 1 mM DTT, 0.25 mM spermidine) supplemented with 0.5% Triton X-100 and protease inhibitors (final concentrations of 2 μg/ml leupeptin, 5 μg/ml aprotinin, 1 mM phenylmethylsulfonyl fluoride, and 1 μg/ml pepstatin A). Cell lysates were centrifuged at 2000 xg for 10 mins at 4LJC and the supernatant loaded on top of a 15 ml 10-30% SDG prepared in buffer A supplemented with 0.1 mg/ml cycloheximide and protease inhibitors. Cycloheximide was used to prevent polysome runoff [42] SDG was centrifuged at 117,100xg for 3.5 hrs at 4LJC in a Sorvall WX 80+ Ultracentrifuge in a Surespin 630 rotor in 17 ml tubes (Thermo, 79386). Fractions (500 ul) were collected from top to bottom of the gradient (30 fractions). SLFN14 was detected by immunoblotting with an anti-FLAG antibody in each fraction, loading per lane 12 ul; whereas RNA was extracted from 400 ul of the fraction with TRIzol LS reagent, as described above.

### *In silico* analysis

Codon adaptation index was determined with the CAIcal program [http://genomes.urv.cat/CAIcal [73]], as described in [74] using as reference the human codon usage table (http://genomes.urv.cat/CAIcal/CU_human_nature). SLFN14 molecular weight was predicted using Expasy. Biorender software was used to generate the graphical abstract.

### Statistical Analysis

GraphPad Prism version 9.4.1 was used for statistical analysis. One-way ANOVA was used to test the impact of human and mouse SLFN14 on the expression of the proteins of interest, and the Dunnett’s *post hoc* test was used to identify significant differences between cells expressing empty plasmid (control group) and cells expressing SLFN14 proteins (experimental groups). Two-tailed, two-sample t test was used to evaluate the statistically significant of experiments with only two groups (control and experimental). Experiments where the comparison was between a specific control and a specific experimental group one-way ANOVA and Bonferroni *post hoc* test was utilized. p-values were indicated as follow: no significant (ns) > 0.05, * ≤ 0.05, ** ≤ 0.01, *** ≤ 0.001, **** ≤ 0.0001.

#### Accession numbers

The sequences of the 5’end of the SLFN14 exon 3 that we detected in MOLT-3 cells is deposited under OP548624 and OP548623.

## Abbreviations

(SLFN): Schlafen
(CAI): codon adaptation index
(MFI): Mean Fluorescence Intensity
(RIN^e^): RNA integrity numbers equivalent
(Ct): PCR cycle threshold
(HRP): horseradish peroxidase

## ACKNOLEDGMENTS

We thank Dr. Georgialina Rodriguez (University of Texas at EL Paso, UTEP) for facilitating access to the electroporator, Dr. Armando Varela (UTEP) for training in using the confocal microscope, Ana Betancourt (UTEP) for running the Agilent Technologies 4200 TapeStation analysis, and Dr. David N. Levy (NYU) for sharing the HIV-1 expression plasmid pNLENG1-ES-IRES. Plasmid pARP-8675, contributed by Drs. Yingying Li, Feng Gao and Beatrice H. Hahn, was obtained through the NIH HIV Reagent Program, Division of AIDS, NIAID, NIH.

## FUNDING

This research was funded by the National Institute on Minority Health and Health Disparities (NIMHD), a component of the National Institutes of Health (NIH) (5U54MD007592). Genomic Analysis Core, the Cytometry Screening and Imaging Core, and the Cell Characterization and Biorepository Core were supported by Research Centers in Minority Institutions Program (5G12MD007592 and 5U54MD007592) from the NIMHD, NIH, to the Border Biomedical Research Center at the University of Texas at EL Paso.

---

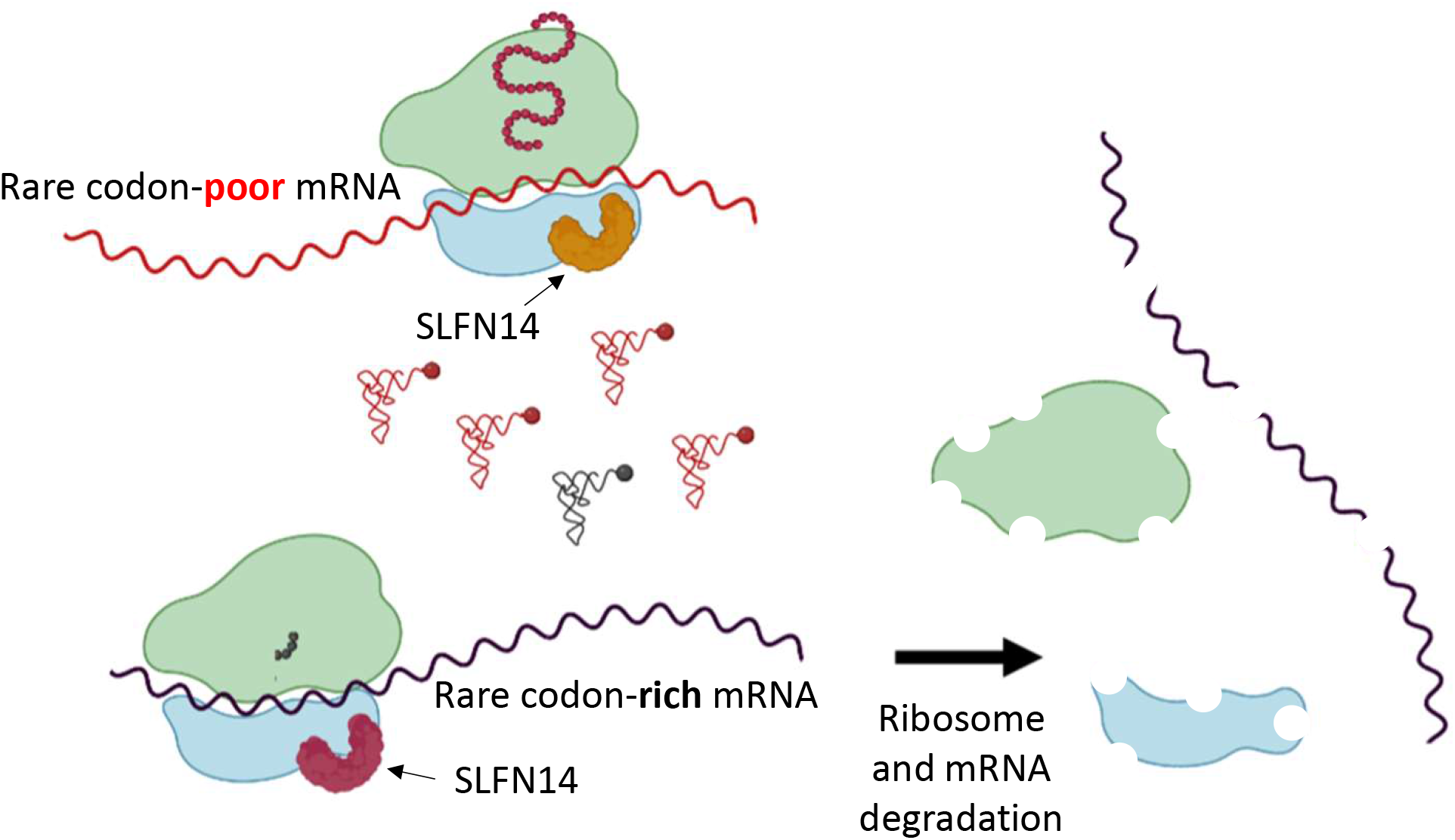

## Notes

### Competing Interest Statement

The authors have declared no competing interest.

